# Forward and backward blocking in statistical learning

**DOI:** 10.1101/2022.02.07.479428

**Authors:** İlayda Nazlı, Ambra Ferrari, Christoph Huber-Huber, Floris P. de Lange

## Abstract

Prediction errors have a prominent role in many forms of learning. For example, in reinforcement learning, agents learn by updating the association between states and outcomes as a function of the prediction error elicited by the event. One paradigm often used to study error-driven learning is blocking. In forward blocking, participants are first presented with stimulus A, followed by outcome X (A→X). In the second phase, A and B are presented together, followed by X (AB→X). Here, A→X blocks the formation of B→X, given that X is already fully predicted by A. In backward blocking, the order of phases is reversed. Here, the association between B and X that is formed during the first learning phase of AB→X is weakened when participants learn exclusively A→X in the second phase. The present study asked the question whether forward and backward blocking occur during visual statistical learning, the largely automatic and incidental learning of the statistical structure of the environment. In a series of studies, using both forward and backward blocking, we observed robust statistical learning of temporal associations among pairs of images. While we found no compelling evidence for forward blocking, we observed reliable backward blocking in visual statistical learning.

## Forward and backward blocking in statistical learning

Learning is an essential feat of animal cognition. It allows us to build and refine our internal models of the world, so that we predict and flexibly adapt to our dynamic environment. A key feature of learning is the ability to form associations between events that take place in a systematic relationship across space or time (Gershman, 2017). For example, in a typical classical conditioning experiment (Pavlov, 1927), a dog automatically salivates (i.e., unconditioned response) in response to food (i.e., outcome or unconditioned stimulus). During conditioning, the sound of a bell (i.e., cue or conditioned stimulus) is repeatedly paired with the food. Once conditioning is accomplished, the bell itself elicits salivation (i.e., conditioned response).

Cue competition is a crucial category of phenomena in associative learning. It refers to the observation that learning which cues predict an outcome not only depends on the presence of the cues before the outcome. Rather, cues compete with each other to gain predictive power over the outcome, and this moderates the learning process (Boddez et al., 2014; De Houwer et al., 2005; Luque et al., 2018; Schmidt & De Houwer, 2019).

One key example of cue competition is Kamin blocking, also known as forward blocking (Kamin, 1969). In a typical forward blocking paradigm (see Table 1), observers first learn the association between cue A and outcome X (A→X), and later they are trained with the association between cues A + B and outcome X (AB→X). As a result of forward blocking, observers learn the association between cue B and outcome X less strongly, because X is already completely predicted by cue A. In other words, the previously learned A-X association blocks learning the association between cue B and outcome X. Forward blocking cannot be explained by simple contiguity-dependent Hebbian associative learning (Hebb, 1949). Thereby, it suggests that the simple temporal co-occurrence of different stimuli is not sufficient for learning to occur. Instead, the model developed by Rescorla and Wagner (1972) provides an explanation for blocking (though see Spicer et al., 2021 for a modification of the traditional model). According to the Rescorla-Wagner model, changes in associative strength are determined by the amount of discrepancy between the expected and the observed outcome, i.e. the prediction error. In the forward blocking procedure, the previously learned A→X association prevents the formation of an associative link between the second cue B and the outcome X, because the cue A already minimizes the prediction error during the exposure to the A→X pairs in the first training phase.

**Table 1.** General experimental design.

|  | *Training phase 1* | *Training phase 2* | *Test phase* |
| --- | --- | --- | --- |
| Forward blocking | $A \rightarrow X$ | $AB \rightarrow X$ | $A \rightarrow X$ |
| | | $CD \rightarrow Y$ | $B \rightarrow X$ |
| | | | $D \rightarrow Y$ |
| Backward blocking | $AB \rightarrow X$ | $A \rightarrow X$ | $A \rightarrow X$ |
| | $CD \rightarrow Y$ | | $B \rightarrow X$ |
| | | | $D \rightarrow Y$ |

Note. Letters denote conditions (i.e., A for Antedating, B for Blocked, C-D for Control).

A similar, but distinct form of cue competition is backward blocking, which is an example of retrospective revaluation: a change in the associative strength occurs because the association between the companion cue (i.e., the cue that is previously associated with the target cue and outcome) and the outcome is revaluated. In the backward blocking paradigm (Shanks, 1985), observers are first trained with AB→X association, and subsequently with A→X association. In spite of the reversed order of training phases compared to forward blocking, backward blocking leads to a similar outcome as forward blocking: a lack of association between blocked cue B and outcome X. Here, in the first training phase, both A-X and B-X associations are formed equally (i.e., depending on the saliency of cues). However, in the second training phase, as observers are trained with A→X association, the associative strength between cue A and outcome X becomes stronger, which in turn weakens the association between cue B and outcome X. While this form of retrospective revaluation cannot be explained by the traditional Rescorla – Wagner model, as this model assumes that the relevant cue must be present in order to change the associative strength (Kruschke, 2008; Miller & Witnauer, 2016; Rescorla & Wagner, 1972), backward blocking can be successfully modeled by a slightly revised version of the traditional model. For example, backward blocking can be explained by a Rescorla-Wagner learning model that assigns non-zero salience to non-presented blocked stimuli whose memories or representations are retrieved by competing stimuli that had previously been paired with those blocked stimuli (Van Hamme & Wasserman, 1994) or by a Bayesian generalization of the Rescorla – Wagner model, the Kalman filter (Gershman, 2015; Kruschke, 2008), where the weights of all possible cues are updated simultaneously, and the sum of all possible weights equals to 1.

In typical blocking experiments, associations are learned either when the outcome is a reward (Aggarwal et al., 2020; Aggarwal & Wickens, 2020; Sharpe et al., 2017; Steinberg et al., 2013) or when performance-related feedback is provided (Blanco et al., 2014; Kruschke & Blair, 2000; Le Pelley et al., 2005, 2007; Luque et al., 2018; Mitchell et al., 2005, 2006). This provides support that reinforcement learning (i.e., learning associations between events via trial and error) relies on an error-driven learning algorithm (Gershman & Daw, 2017). Another powerful form of learning is known as statistical learning, often defined as the automatic and incidental extraction of regularities from the environment (Batterink et al., 2019; Frost et al., 2019; Saffran et al., 1996; Sherman et al., 2020; Turk-Browne et al., 2010). In the context of statistical learning, we have limited information about how the learning process itself occurs. Several studies are suggestive of the fact that statistical learning may indeed similarly rely on prediction errors. In rats, dopaminergic activity in the ventral tegmental area is important for the formation of an association between two non-rewarding stimuli (Keiflin et al., 2019; Sharpe et al., 2017). In humans, statistical learning involves the ventral striatum (Klein-Flügge et al., 2019), which has been hypothesized to signal prediction errors (Klein-Flügge et al., 2019; McClure et al., 2003; O’Doherty et al., 2004).

However, other researchers, using variants of forward blocking, did not find clear-cut evidence for error-driven statistical learning. Beesley and Shanks (2012) did not observe any forward blocking in contextual cueing experiments, where participants incidentally learnt the spatial relationship among distractors and targets in a visual search task. This procedure however deviates from classic forward blocking paradigms, which rely on a *temporal* prediction between a cue and a future outcome (Aggarwal et al., 2020; Aggarwal & Wickens, 2020; Blanco et al., 2014; De Houwer et al., 2005; De Houwer & Beckers, 2003; Kruschke & Blair, 2000; Le Pelley et al., 2005, 2007; Luque et al., 2018; Mitchell et al., 2006; Steinberg et al., 2013; Vandorpe et al., 2005). Two subsequent experiments (Morís et al., 2014; Schmidt & De Houwer, 2019) observed forward blocking of temporal associations only for material that was intentionally learnt, but not for incidentally learnt stimulus associations. Such learning conditions substantially deviate from a typical statistical learning scenario, where observers automatically extract regularities without intention or awareness (Batterink et al., 2019; Frost et al., 2019; Sherman et al., 2020; Turk-Browne et al., 2010). Despite few studies investigating forward blocking in incidental learning (Blanco et al., 2014; Kruschke & Blair, 2000; Le Pelley et al., 2005, 2007; Mitchell et al., 2006), to the best of our knowledge, no study has examined backward blocking in incidental learning. According to the attentional account model (Mackintosh, 1975), forward blocking occurs as a result of less attention devoted to the blocked cue. Namely, in the first training phase observers learn that cue A predicts outcome X and cue A therefore becomes a relevant and attended cue. Therefore, in the second training phase where novel cue B is presented together with cue A, more attention is paid to the relevant and predictive cue A, compared to the novel predictive cue B. The lower attention to cue B leads to the failure to associate cue B with outcome X. This attentional explanation would however not be able to explain backward blocking, given that equal attention is paid to cues A and B in the first phase. Hence, the use of backward blocking enables us to examine more closely the attentional account of blocking and, crucially, test for retrospective revaluation in statistical learning.

Therefore, we set out to examine forward and backward blocking during statistical learning in a series of experiments. In some statistical learning experiments, participants are exposed to a continuous stream of stimuli containing statistical regularities (Batterink et al., 2019; Batterink & Paller, 2017; Henin et al., 2021; Saffran et al., 1996; Turk-Browne et al., 2005, 2009). Other studies have instead presented two successive stimuli on each trial, with conditional probabilities controlling their pairing (Richter et al., 2018; Richter & de Lange, 2019). In terms of neural processing, both continuous streams (Kaposvari et al., 2018) and pairs (Meyer & Olson, 2011) show identical modulations of sensory responses after statistical learning, suggesting that both paradigms elicit similar learning processes. We opted for pairs of stimuli in order to connect our study to the classic forward and backward blocking paradigms (Kamin, 1969; Shanks, 1985). On every trial, we presented participants with two consecutive visual object stimuli and asked them to categorize the trailing object as either electronic or non-electronic. Unbeknownst to participants, we manipulated the conditional probabilities between the leading and trailing stimuli, such that each trailing image could be predicted on the basis of its preceding, leading image. After learning, we evaluated statistical learning by presenting participants with expected and unexpected image pairs and measuring their reaction time for categorization judgments of the trailing image. Successful learning was indexed by faster reaction times to expected relative to unexpected trailing stimuli (Hunt & Aslin, 2001; Richter & de Lange, 2019; Turk-Browne et al., 2005).

## Experiment 1

## Method

### Preregistration and data availability

All experiments were preregistered on the Open Science Framework (https://osf.io/r243e for Experiment 1; https://osf.io/7kmtv for Experiment 2). All data and code used for the analyses are freely available on the Donders Repository (for review only -https://data.donders.ru.nl/login/reviewer-183657338/X8JOMneRBVvc26TYCJEg9xS5uNl-PvbY2l-48Ft5Vuk). Deviations from the preregistration are mentioned as such and justified in the corresponding sections below.

### Participants

The experiment was performed online by using the Gorilla platform (Anwyl-Irvine et al., 2020), and participants were recruited through the Prolific platform (https://www.prolific.co/). 92 participants performed the experiment. 42 of them were excluded before they finished the experiment based on a priori exclusion criteria (see section ‘Exclusion and inclusion criteria’ above). As a result, fifty participants (18 females; mean age 25.80, range 18-40 years) were included in the data analysis. This final number of included participants was derived from the following a priori power calculation: we aimed for 90% power to detect a medium effect size (Cohen’s *d*_z_ = 0.5), as derived from a Supplementary Experiment 1 (N=100).

All participants had normal or corrected to normal vision, normal hearing and no history of neurological or psychiatric conditions. They provided written informed consent and received financial reimbursement (8 euro per hour) for their participation in the experiment. The study followed the guidelines for ethical treatment of research participants by CMO 2014/288 region Arnhem-Nijmegen, The Netherlands.

### Experimental design

In each experimental trial, participants were exposed to two images presented on the left or right side of the central fixation point in quick succession: a leading stimulus was followed by a trailing stimulus. For each participant, there were 4 leading objects and 4 trailing stimuli objects. Everyday objects were randomly chosen from a pool of 64 stimuli derived from Brady et al. (2008) per participant, thereby eliminating potential effects induced by individual image features at the group level. In each stimulus set, 50% of objects were electronic (consisting of electronic components and/or requiring electricity to function) and 50% were non-electronic. The expectation manipulation consisted of a repeated pairing of objects in which the leading object predicted the identity of the trailing object, thus over time making the trailing object expected given the leading object. Importantly, each trailing object was only (un)expected depending on which leading object it was preceded by. Thus, each trailing object served both as an expected and unexpected object depending on the leading object at test phase. In addition, trial order was pseudo-randomized, with the pairs distributed equally over time. In sum, any difference between expected and unexpected occurrences cannot be explained in terms of familiarity, adaptation, or trial history. In addition, object position (left / right) was counterbalanced with respect to Expectation (expected / unexpected) and Condition (antedating / blocked / control). In other words, leading and trailing objects appeared equally often on the left or right side of the central fixation point across trials. As a result, the expectation manipulation did not depend on spatial position. Also, both hemi-fields were equally task-relevant, which fostered participants’ attention to both sides. Throughout the experiment, participants needed to categorize the trailing object as electronic or non-electronic as fast as possible. This task was aimed at assessing any implicit reaction time (RT) benefits due to incidental learning of the temporal statistical regularities: upon learning, leading object could be used to predict the correct categorization response before the trailing object appeared. In addition to the main object categorization task, there was an oddball detection task involving the leading stimuli in the training phases (16% of all trials per participant): participants were required to press a specific button as soon as they saw an animate leading stimulus. The aim of the animate detection task was to ensure that participants also paid attention to the leading stimuli, such that the association would be better learnt. For each participant, 4 animate leading stimuli (i.e., 2 for antedating leading stimulus and 2 for blocked leading stimulus) were randomly chosen from a pool of 8 stimuli (Brady et al., 2008). Finally, there were attention check trials where participants were simply asked to press a specific key based on a message on screen (e.g., "Press left-arrow key"). The aim of these trials (7% of all trials per participant) was to monitor participants’ vigilance (see ‘Exclusion and inclusion criteria’). A fixation bull’s-eye was presented in the center of the screen throughout the experiment.

The blocking paradigm comprised two consecutive training phases, followed by one test phase (see Figure 1a). During the two training phases, leading objects were perfectly predictive of their respective trailing objects (i.e. P(trailing | leading = 1) ; see Figure 1b). Participants were not informed about this deterministic association, nor were they instructed to learn this association at the beginning of the experiment. Therefore, the pair associations were likely learned incidentally.

**Figure 1.**
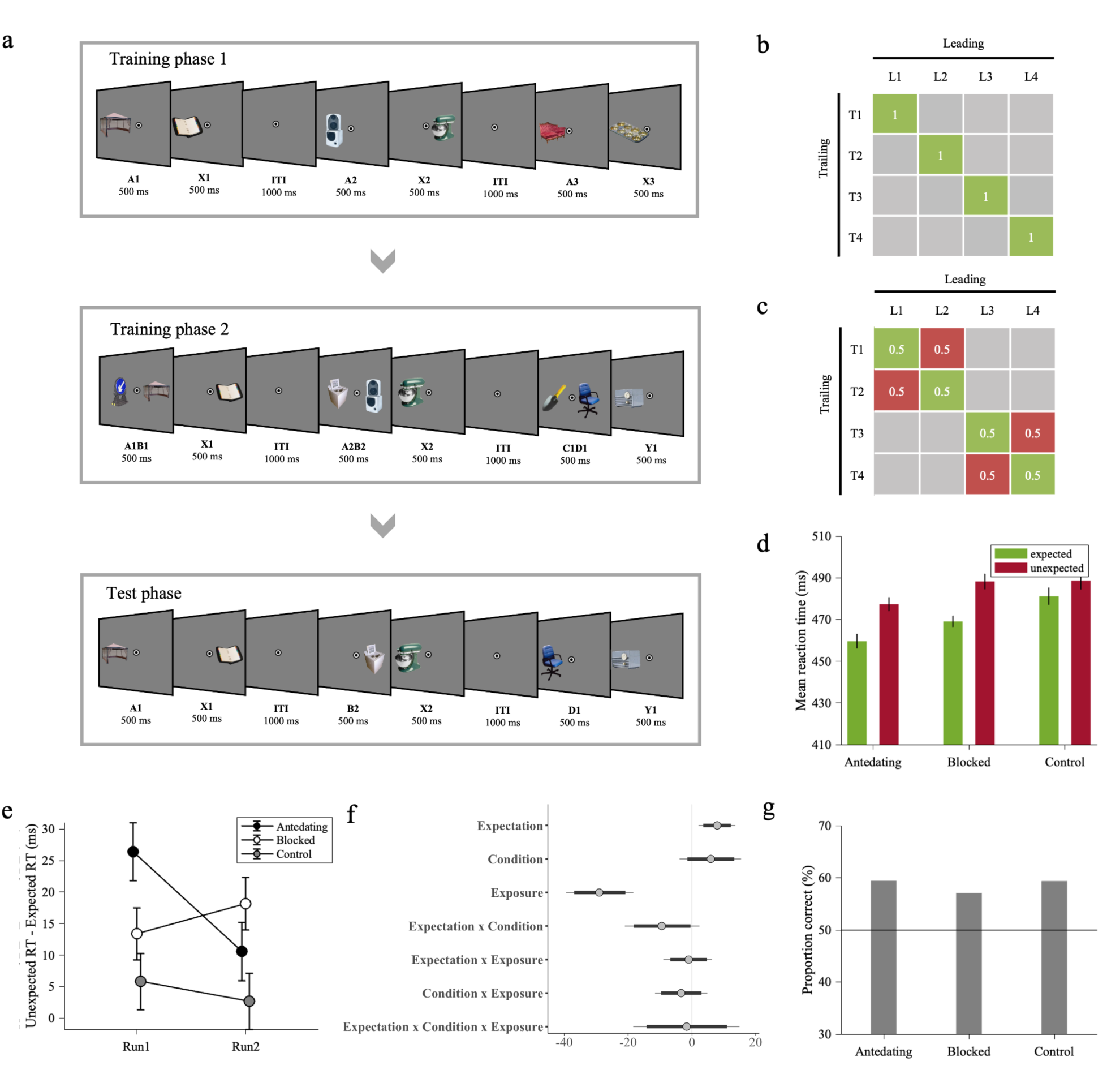
Experimental procedure and results of Experiment 1. *Note.* (a) Experiment 1 comprised two training phases (training phase 1 and training phase 2) and a test phase. On every trial throughout the experiment, participants saw a pair of consecutively presented stimuli, i.e., a leading object followed by a trailing object. In training phase 1, the antedating leading object (i.e., A) was followed by a specific trailing object. In training phase 2, a novel blocked leading object (i.e., B) was presented in compound, along with the antedating (A) leading object (i.e., AB), and followed by the same trailing object from the antedating stimulus in training phase 1. In addition, we introduced novel control compound leading (i.e., CD) and trailing (i.e., Y) objects. In the test phase, antedating, blocked or control leading stimuli were followed by the associated (expected) or not associated (unexpected) trailing object. There were four different object pairs for AB→X and CD→Y. Throughout the experiment, participants performed a categorization task on the trailing object. They reported, as fast as possible, whether the trailing object was electronic or non-electronic. (b) Statistical regularities depicted as image transition matrix with stimuli pairs in training phase 1 and training phase 2. Ls represent leading stimuli, and Ts represent trailing stimuli. There were 16 different leading objects and 8 different trailing objects coming from four different AB→X and CD→Y pairs. (c) Statistical regularities depicted as image transition matrix with stimuli pairs in test phase. Green cells represent expected pairs, and red cells represent unexpected pairs. (d) Across participants’ mean reaction times as a function of Expectation (expected / unexpected) and Condition (antedating / blocked / control). Reaction times were faster to expected than unexpected trailing objects in each condition. The reaction time difference between expected and unexpected trials was greater in blocked than control trials, providing evidence for the absence of blocking effect and the augmentation of learning. (e) Across participants’ mean reaction time difference between expected and unexpected trials as a function of time. Please note that we split data into successive runs for visualization purposes only; data analysis was performed with number of trials as a continuous fixed factor (Exposure). The decrease in reaction time difference between expected and unexpected trials over exposure showed rapid extinction in learning antedating condition. (f) Posterior coefficient estimates of effects of the model jointly analyzing blocked and control conditions with error bars representing 95% confidence intervals. Estimates indicate significant results when they do not overlap with zero. (g) Across participants’ proportion correct responses in pair recognition test. Participants showed slightly above chance-level performance in all conditions indicating whether the trailing object was likely or unlikely given the leading object.

Note that the participants may, however, still develop explicit knowledge of the associations over the course of the experiment, which we tested in a final recognition task. In training phase 1, the leading object (A) was always followed by the same trailing object (X). In training phase 2, a novel leading object (blocked [B] leading object) was presented along with the leading object presented in training phase 1 (antedating [A] leading stimulus), hence creating a compound stimulus (AB). This was followed by the same trailing object (X) as in training phase 1. In addition, two novel leading (object + object [CD]) and a trailing (object [Y]) objects were presented as a control condition. In the test phase, the leading stimulus of each condition (antedating [A] / blocked [B] / control [D]) was presented alone, followed by either the expected trailing object (based on the training phases), or an unexpected trailing object. Expected and unexpected object pairs were presented equally often to prevent any learning at this final test stage (see Figure 1c). In the test phase, control (D) trials were compared to blocked (B) trials to assess blocking while controlling for the amount of exposure. It should be noted that the amount of exposure to trailing object X and trailing object Y are not the same, given that trailing object Y was only introduced in the second learning phase. This difference is an inevitable feature in classic blocking paradigms. If we would have presented trailing object Y in isolation in an additional experimental phase or if we would have paired Y with another leading stimulus in training phase 1, this could have elicited latent inhibition (i.e., difficulty in learning associations as a result of pre-exposure, McLaren & Mackintosh, 2000). Thus, we opted for the classic blocking paradigm. Furthermore, the control trials in the test phase allowed us to assess whether new associations were learned during training phase 2.

Data was collected during one single session per participant. Firstly, participants familiarized themselves with all trailing objects (both X and Y). In each trial, an object image was presented for 3500 ms, and participants had 1500 ms to categorize the object image as electronic or non-electronic (via a keyboard key press, keys counterbalanced across participants). Then, written feedback indicated the true category and the name of the object for 2000 ms (8 pairs × 2 trials / pairs = 16 trials in total). Afterwards, participants performed the experiment (i.e., training phase 1, training phase 2 and test phase). In each trial, the leading and trailing objects were presented for 500 ms successively with no inter-stimulus interval, followed by a 1500 ms inter-trial interval.

Participants categorized the trailing object as electronic or non-electronic as fast as possible (via keyboard key press, keys counterbalanced across participants). Training phase 1 and training phase 2 started with a short practice period (practice training phase 1: 4 pairs × 4 trials / pairs = 16 trials in total; practice training phase 2: 8 pairs × 4 trials / pairs = 32 trials in total). After each practice, participants completed the training phases (training phase 1: 4 object pairs × 30 trials = 120 trials in total; training phase 2: 8 object pairs × 30 trials = 240 trials in total). In addition, animate detection and attention check trials (see above) were pseudo-randomly interspersed throughout the training phases without repetitions in successive trials. Afterwards, participants completed the test phase (12 pairs × 16 trials = 180 trials in total). Crucially, for each leading object, both expected and unexpected trailing objects belonged to the same category (electronic or non-electronic). This ensured that differences in RTs during object categorization would not arise by mere response adjustments costs, but instead reflected perceptual surprise to unexpected trailing objects.

Finally, at the end of the experiment participants performed a pair recognition task to probe their explicit knowledge of the statistical regularities. Before starting the recognition task, participants were informed about the presence of statistical regularities among leading and trailing images in the previous experimental phases (i.e., training phases 1 and 2), and they were asked to indicate whether the trailing object was likely or unlikely given the leading stimulus according to what they saw during these previous phases. Participants familiarized themselves with the procedure via a brief practice (12 pairs × 2 trials / pairs = 24 trials in total) before completing the recognition task (12 pairs × 8 trials / pairs = 96 trials in total).

### Exclusion and inclusion criteria

The online experiment was terminated if the percentage of correct responses during object categorization was below 80% (threshold was defined based on a preliminary pilot study) in any training or test phase (see ‘Experimental design’ and Figure 1a) or if the percentage of correct responses in attention check trials was below 80% in any of the experimental phases (see section ‘Experimental design’).

Prior to the main data analysis, we discarded trials with no responses, wrong responses, or anticipated responses (i.e., response time < 200 ms). We also rejected trial outliers (response times exceeding 3 MAD from mean RT of each participant) and subject outliers (participants whose RTs exceeded 3 MAD from the group mean). For the accuracy analysis of the pair recognition task, we rejected trial outliers in terms of response speed (response times exceeding 3 MAD from mean RT of each participant).

### Data analysis

We analyzed the RT data in the test phase in order to test for incidental learning of predictable stimulus transitions: upon learning, participants were hypothesized to react faster to expected relative to unexpected trailing stimuli (Richter et al., 2018, Richter & de Lange, 2019). We did not statistically analyze the accuracy data in the test phase, given that the categorization task was not challenging, and performance was near ceiling levels (97% in Experiment 1 and 97% in Experiment 2). Furthermore, we analyzed the accuracy data in the pair recognition test to assess participants’ explicit knowledge about learnt statistical regularities. For both analyses, we used a Bayesian mixed effect model approach. The Bayesian framework allows a three-way distinction between evidence for an effect, evidence for no effect, and absence of evidence (Dienes, 2016; Keysers et al., 2020).

This three-way distinction is important in the present study because it allowed us to draw conclusions from the initial experiment, consider alternative explanations, and run follow-up experiments to test these alternative explanations. An additional reason for this approach was the violation of the normality assumption for repeated measures ANOVAs of response times. Data were analyzed using the *brm* function of the BRMS package (Bürkner, 2017) in R. In the Supplementary information, we additionally provide classic frequentist analyses (i.e., ANCOVA of the reaction time data of the test phase and one-way ANOVA of the accuracy data of the pair recognition test) for comparability with previous studies and to verify that our conclusions do not depend on the analytical framework employed. Furthermore, in supplementary tables we provide post-hoc Bayesian mixed effect models that follow significant interaction effects.

#### Analysis of RT data in test phase

Firstly, we modeled the behavioral data of the antedating condition, where one leading stimulus was followed by one trailing stimulus. This served as a sanity check to verify the baseline assumption that participants were able to learn the temporal association between the leading and trailing stimuli. The model of the antedating (A) condition included reaction time as dependent variable and Expectation (unexpected / expected) as a fixed factor. To model the overall effect of time on task, we included Exposure as a continuous numeric predictor.

Exposure was scaled between -1 and 1 to be numerically in the same range as the other factors, which aids model convergence. For the interpretation of the results, the model coefficient for Exposure represents the increase in RT from the first to the last exposure. Finally, we included the interaction between Exposure and Expectation in the model, to probe extinction of the learnt associations. Namely, during the test phase participants were exposed equally often to expected and unexpected stimulus pairs, potentially resulting in extinction of the RT advantage for expected stimuli over time. The model included a full random effect structure (i.e., a random intercept and slopes for all within-participant effects).

Secondly, we determined whether there was blocking by jointly modeling the blocked (B) and control (D) conditions. The model of blocked and control conditions included reaction time as a dependent variable and Expectation (unexpected / expected), Condition (control / blocked) and Exposure as fixed independent variables. We included the interaction between Expectation and Condition to test for the blocking effect. The contrasts of the factors Expectation and Condition were coded as successive difference contrasts. Exposure was a continuous predictor scaled between -1 and 1, as in the antedating condition analysis. Again, we also modeled extinction (Expectation × Exposure interaction) and its interaction with Condition to probe for potential differences in extinction between conditions. We adjusted the priors of the main effect of Expectation and Exposure and the prior of their interaction based on the posteriors of pilot experiments. Each prior was centered according to the median of the respective posterior estimate, and its standard deviation equated to the posterior estimate error times two to make the priors less informative. The prior for the Condition effect and its interaction with Expectation, i.e., blocking effect, was centered at zero. Note that specifying the priors in this way turns the estimates of Expectation and Exposure effects of Experiment 1 into the combined evidence from pilot experiments *and* Experiment *1*. Crucially, the pattern of results from Experiment 1 was exactly the same when not only the priors for the Condition effect but also for Expectation and Exposure were centered at zero. Further details and the complete model parametrization can be found in the R codes provided on the Donders Repository. The response time data was modelled using the ex-gaussian family and four chains with 25,000 iterations each (12,500 warm up) per chain and inspected for chain convergence. We report posterior fixed effects model coefficients. Coefficients were accepted as convincing statistical evidence, analogously to statistically significant in a frequentist framework, if the associated 95% posterior credible intervals were non-overlapping with zero.

#### Analyses of accuracy data in pair recognition test

Firstly, we determined whether accuracy was above chance level within each condition (antedating / blocked / control). Hence, we created three separate binomial mixed-effects models with response error as dependent variable. If accuracy was above chance level within each condition, we then determined whether there was a blocking effect in the explicit knowledge of implicitly learned associations. To do so, we created a binomial mixed-effects model with response error as binary dependent variable and Condition (blocked / control) as fixed factor. The models included a full random effect structure (i.e., a random intercept and slopes for the within-participant effects). The models were constructed using weakly informative priors centered at zero. All accuracy models were fit using Bernoulli family and four chains with 25,000 iterations each (12,500 warm up) per chain and inspected for chain convergence. With respect to significance and amount of evidence we used the same criteria as for the RT data.

## Results

### Analyses of RT data in test phase

Firstly, we compared the reaction times of expected and unexpected trials in the antedating condition (see Table 2). We observed faster reaction times in expected (460 ms) than in unexpected (477 ms) trials (b = 10.81, CI = [5.04, 16.16], Cohen’s *d*_z_ = 0.61, see Figure 1d), indicating successful learning of conditional probabilities and the consequent behavioral benefit of expectation in terms of response speed. In addition, we evaluated how this learning effect changed across exposure. Again, we observed an interaction effect between expectation and exposure (b = -9.01, CI = [-16.83, -1.18]), indicating that learning showed rapid extinction (expectation effect for run 1: 26 ms, run 2: 11 ms; see Figure 1e).

**Table 2.** Posterior fixed effects of the model of antedating condition on reaction times in Experiment 1. Estimate, estimation error, lower/upper limit of 95% profile credible intervals.

| Predictors | *Estimate* | *Est. Error* | *CI (95%)* |
| --- | --- | --- | --- |
| Intercept | 474.88 | 10.37 | 454.81 – 495.21 |
| Expectation | 10.81 | 2.83 | 5.04 – 16.16 |
| Exposure | -23.39 | 3.70 | -30.63 – -16.08 |
| Expectation × Exposure | -9.01 | 4.00 | -16.83 – -1.18 |

Next, we modeled the blocked and control conditions to test whether we found blocking (see Table 3 and Figure 1f). There was an interaction effect between expectation and condition (b = - 9.48, CI = [-18.26, -0.45], Cohen’s *d*_z_ = -0.26, see Figure 1b. We performed separate analyses for the blocked and control conditions to test for the presence of an expectation effect in each condition respectively. The reaction times in expected (481 ms) and unexpected (489) trials were not different from each other in the control condition (b = 4.36, CI = [-0.73, 9.51], Cohen’s *d*_z_ = 0.20, see Table S1). On the other hand, reaction times were clearly faster in expected (469 ms) than in unexpected (488 ms) trials of the blocked condition (b = 10.11, CI = [4.82, 15.16], Cohen’s *d*_z_ = 0.65, see Table S2). Interestingly, this is exactly the opposite pattern of what would be expected under blocking, and rather supports better learning of the associations among blocked stimuli than control stimuli. Extinction was not different between blocked and control conditions (b = -1.63, CI = [-14.19, 11.00]; expectation effect in blocked condition for run 1: 13 ms, run 2: 18 ms; expectation effect in control condition for run 1: 6 ms, run 2: 3 ms; see Figure 1c).

**Table 3.**
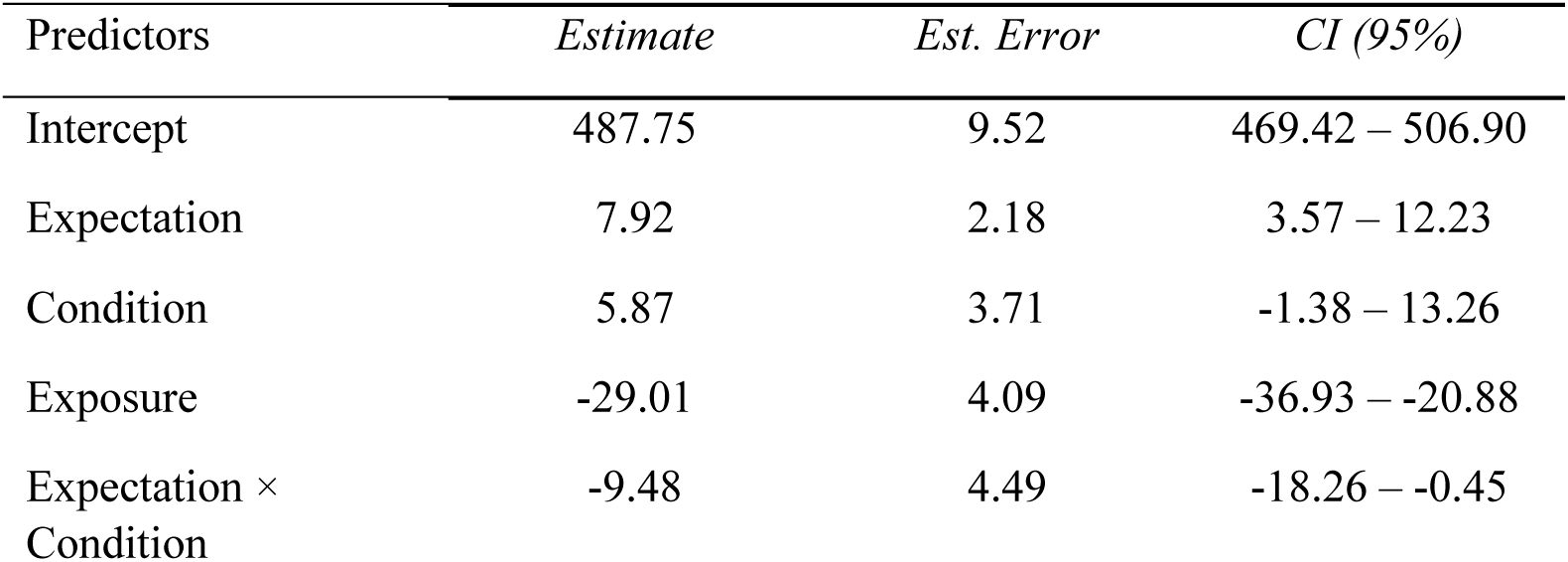
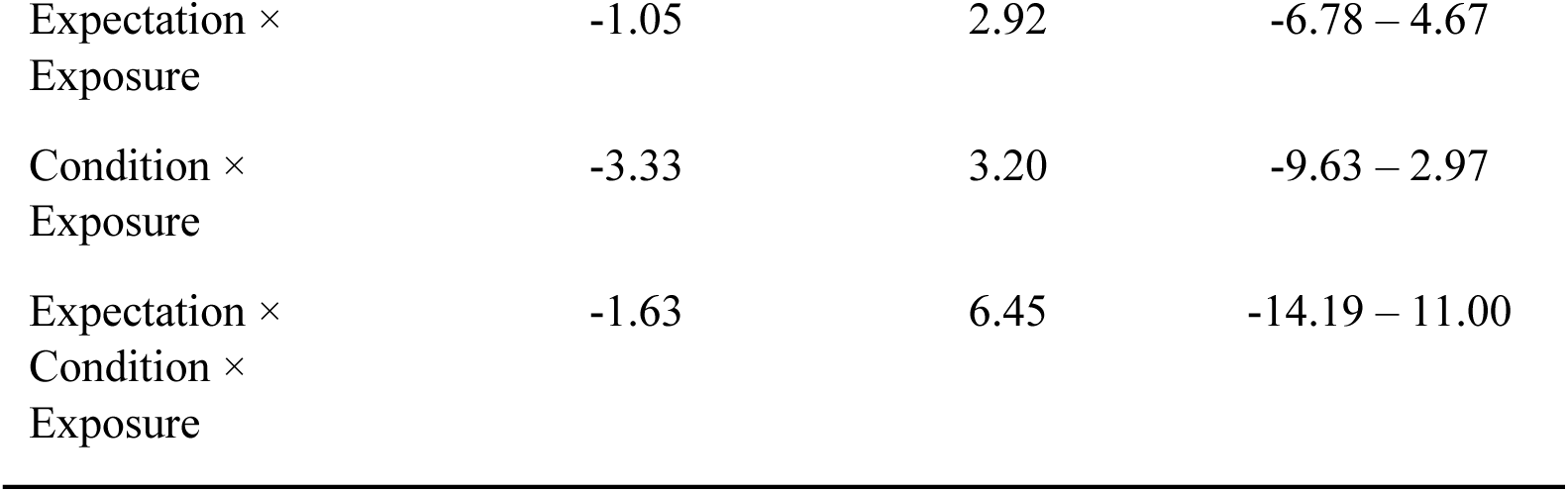
Posterior fixed effects of the model of blocked and control conditions on reaction times in Experiment 1. Estimate, estimation error, lower/upper limit of 95% profile credible intervals.

| Predictors | *Estimate* | *Est. Error* | *CI (95%)* |
| --- | --- | --- | --- |
| Intercept | 487.75 | 9.52 | 469.42 – 506.90 |
| Expectation | 7.92 | 2.18 | 3.57 – 12.23 |
| Condition | 5.87 | 3.71 | -1.38 – 13.26 |
| Exposure | -29.01 | 4.09 | -36.93 – -20.88 |
| Expectation × Condition | -9.48 | 4.49 | -18.26 – -0.45 |
| Expectation ×<br>Exposure | -1.05 | 2.92 | -6.78 – 4.67 |
| Condition ×<br>Exposure | -3.33 | 3.20 | -9.63 – 2.97 |
| Expectation ×<br>Condition ×<br>Exposure | -1.63 | 6.45 | -14.19 – 11.00 |

### Analysis of accuracy data in pair recognition test

Participants showed slightly above-chance level performance in indicating whether the trailing object was likely or unlikely given the leading object in the antedating (proportion correct = 59%; b = 0.39, CI = [0.26, 0.51]), blocked (proportion correct = 57%; b = 0.29, CI = [0.17, 0.42]) and control (proportion correct = 59%; b = 0.39, CI = [0.24, 0.54]) conditions (see Figure 1g). Response errors did not differ between the blocked and control conditions (b = -0.1, CI = [-0.08, 0.29]), indicating the absence of blocking effect for the explicit knowledge of incidentally learned associations.

## Experiment 2

In Experiment 1, we observed a stronger reaction time benefit for B→X compared to control, indicating successful learning and the absence of forward blocking. We speculated that this pattern of results may be explained by the following process: upon learning the A→X association in the first training phase, attention may have shifted to the novel (and therefore potentially more salient) leading image B during the second training phase, thereby enhancing the learning of the B→X association. Importantly, this attentional mechanism is not at play in the related, but distinct paradigm of backward blocking (Shanks, 1985). Here, the order of training phases is reversed compared to forward blocking. Observers are first trained with AB→X association and presented in a subsequent training phase with the A→X association. As a result, both leading objects A and B are equally novel and salient during the first training phase and therefore should be learnt equally well. Therefore, we reasoned that backward blocking may allow us to study blocking without the potentially confounding factors related to novelty and salience. Crucially, this paradigm also allowed us to test for the first time whether retrospective revaluation takes place during statistical learning.

## Method

### Participants

The experiment was performed online by using the Gorilla platform (Anwyl-Irvine et al., 2020), and participants were recruited through the Prolific platform (https://www.prolific.co/).

Eighty-four participants performed the experiment. Thirty-three of them were excluded before they finished the experiment based on a priori exclusion criteria (see section ‘Exclusion and inclusion criteria’ below). One participant was excluded from the final data analysis due to overall excessively fast responses (i.e., 93% of responses being less than 200 ms). As a result, 50 participants were included in the data analysis, as preregistered. This final number of included participants was based on the same power analysis explained above.

All participants had normal or corrected to normal vision, normal hearing and no history of neurological or psychiatric conditions. They provided written informed consent and received financial reimbursement (8 euro per hour) for their participation in the experiment. The study followed the guidelines for ethical treatment of research participants by CMO 2014/288 region Arnhem-Nijmegen, The Netherlands.

### Experimental design

The design and procedure of Experiment 2 was identical in all respects to Experiment 1, apart from the fact that the order of elemental and compound training phases was reversed (see Table 1 and Figure 2a).

**Figure 2.**
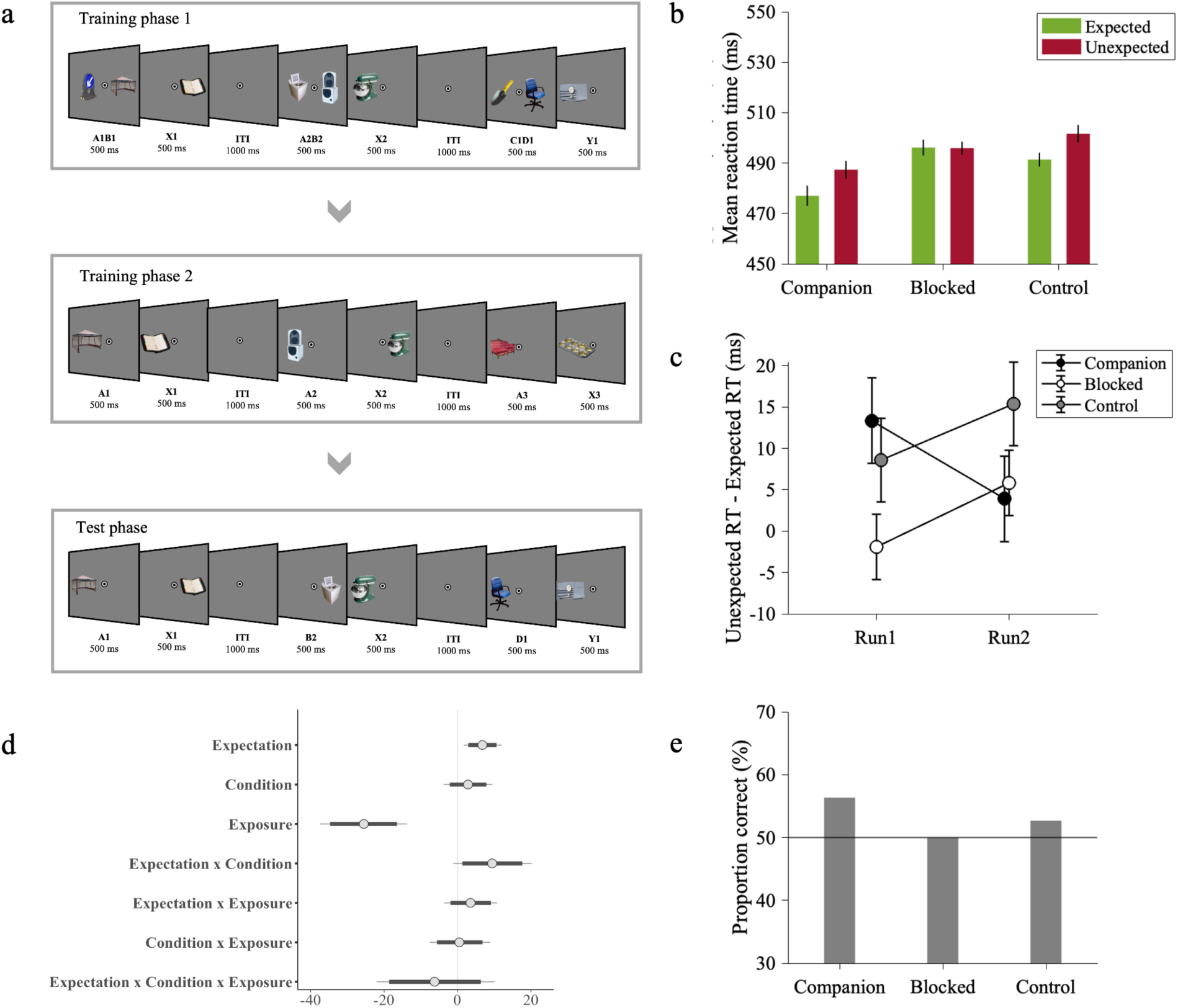
Experimental procedure and results of Experiment 2. *Note.* (a) The design and procedure of Experiment 2 was identical in all respects to Experiment 1, apart from the order of training phases. (b) Across participants’ mean reaction times as a function of Expectation (expected / unexpected) and Condition (companion / blocked / control). Reaction times were faster to expected than unexpected trailing objects in companion and control conditions but not in blocked condition, providing evidence for the presence of backward blocking in statistical learning. (c) Across participants’ mean reaction time difference between expected and unexpected trials as a function of time. There was no extinction in learning in any conditions. (d) Posterior coefficient estimates of effects of the model jointly analyzing blocked and control conditions with error bars representing 95% confidence intervals. Estimates indicate significant results when they do not overlap with zero. (e) Across participants’ proportion correct responses in pair recognition test. Participants showed slightly above chance-level performance in companion condition indicating whether the trailing object was likely or unlikely given the leading object.

### Data analysis

The data analysis of Experiment 2 was the same as for Experiment 1. Also here, we adjusted the priors of the main effect of Expectation and Exposure and the prior of their interaction based on the posteriors of the previous experiment, i.e., Experiment 1, because the stimuli and procedure regarding effects of Expectation and Exposure were exactly the same. Note that specifying the priors in this way turns the results of Experiment 2 with respect to Expectation and Exposure effects into the combined evidence from Experiments 1 *and* 2. Crucially, the pattern of results from Experiment 2 was exactly the same when the priors for Expectation and Exposure were also centered at zero.

### Results

#### Analysis of RT data in test phase

First, we compared the reaction times of expected and unexpected trials in the companion condition to test whether repeated exposure to the pairs of the companion leading object A and trailing object X led to learning their temporal association (see Table 4). We observed faster reaction times in expected (477 ms) than unexpected (487 ms) trials (b = 8.90, CI = [3.53, 14.27], Cohen’s *d*_z_ = 0.35, see Figure 2b), indicating successful learning of stimulus transition probabilities and the consequent behavioral benefit of expectation in terms of response speed. In addition, we tested whether this behavioral benefit remained stable during the test phase or tended to decrease as the exposure increased (i.e., extinction). We did not observe any interaction effect between Expectation and Exposure (b = 4.63, CI = [-12.41, 3.04]), indicating that learning did not show reliable extinction over time (expectation effect for run 1: 13 ms, run 2: 4 ms; see Figure 2c).

**Table 4.** Posterior fixed effects of the model of companion condition on reaction times in Experiment 2. Estimate, estimation error, lower/upper limit of 95% profile credible intervals.

| Predictors | *Estimate* | *Est. Error* | *CI (95%)* |
| --- | --- | --- | --- |
| Intercept | 483.97 | 11.08 | 462.31 – 505.62 |
| Expectation | 8.90 | 2.75 | 3.53 – 14.27 |
| Exposure | -14.12 | 4.38 | -22.81 – -5.66 |
| Expectation ×<br>Exposure | -4.63 | 3.93 | -12.41 – 3.04 |

Next, we moved to our main question and tested for the presence of backward blocking (see Table 5 and Figure 2d). There was an interaction effect between expectation and condition (b = 9.45, CI = [1.34, 17.63], Cohen’s *d*_z_ = 0.32, see Figure 2b). We performed separate analyses for the blocked and control conditions to test for the presence of an expectation effect in each condition respectively. The reaction times were faster in expected (491 ms) than in unexpected (501 ms) trials of the control condition (b = 8.44, CI = [3.60, 13.29], Cohen’s *d*_z_ = 0.39, see Table S3). On the other hand, there was no evidence that reaction times in expected (496 ms) and unexpected (496) trials were different from each other in the blocked condition (b = 2.69, CI = [-2.08, 7.44], Cohen’s *d*_z_ = -0.01, see Table S4). This pattern of results supports the presence of backward blocking. There was no extinction in blocked and control conditions (b = -6.22, CI = [-18.63, 6.38]; expectation effect in blocked condition for run 1: 0 ms, run 2: 6 ms; expectation effect in control condition for run 1: 9 ms, run 2: 15 ms; see Figure 2c).

**Table 5.**
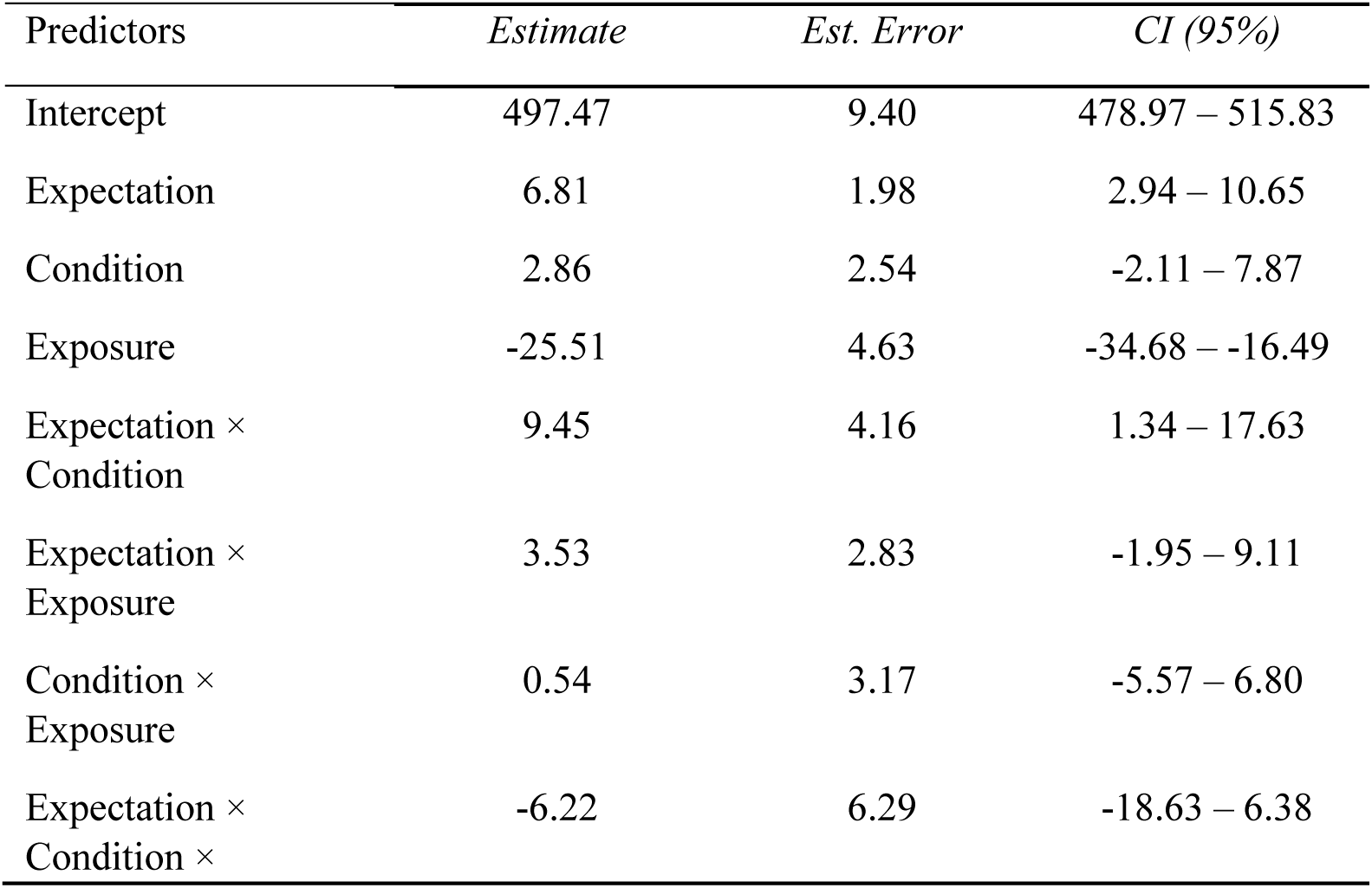
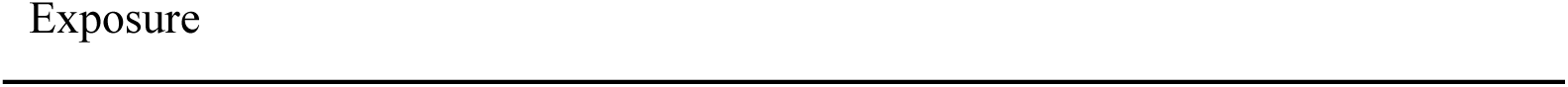
Posterior fixed effects of the model of blocked and control conditions on reaction times in Experiment 2. Estimate, estimation error, lower/upper limit of 95% profile credible intervals.

*Analyses of accuracy data in pair recognition test.* Participants showed slightly above chance-level performance in indicating whether the trailing object was likely or unlikely given the leading stimulus in the companion (proportion correct = 56%; b = 0.25, CI = [0.10, 0.40]), but not in blocked (proportion correct = 50%; b = 0, CI = [-0.11, 0.11]) and control (proportion correct = 53%; b = 0.09, CI = [-0.04, 0.22]) conditions (see Figure 2e).

## Discussion

Statistical learning allows us to detect and learn structure in the environment, with direct benefits for directing our limited processing resources more efficiently to optimize behavior. This results, for example, in more efficient behavioral processing (Fiser & Aslin, 2001, 2002; Hunt & Aslin, 2001; Saffran et al., 1996, 1999) and more efficient neural processing (Batterink & Paller, 2017; Henin et al., 2021; Richter et al., 2018; Richter & de Lange, 2019; Turk-Browne et al., 2009) for predictable than unpredictable events. While the benefits of statistical learning are obvious, the mechanisms of statistical learning itself are less clear. In separate experiments, we used respectively forward and backward blocking (Kamin, 1969; Shanks, 1985) to examine whether cue competition and retrospective revaluation, which have been observed during reinforcement learning, also apply to statistical learning. We found evidence of backward blocking in statistical learning, suggesting a retrospective revaluation process in statistical learning.

In Experiment 1, strong learning of the associations for the blocked (B) stimulus condition; in fact, learning was even stronger for B stimuli compared to control (D) condition, a phenomenon which is sometimes referred to as ‘augmentation’ (Batson & Batsell Jr., 2000; Beesley & Shanks, 2012; Vadillo & Matute, 2010). This pattern of results is opposite to the predictions of forward blocking and provides evidence for the absence of forward blocking in statistical learning. One might argue that overall learning in the antedating and blocked conditions may not have been strong enough to generate forward blocking, given that the reduction in response speed was less than 20 ms. Such a small reaction time difference is, however, common in statistical learning (Richter et. al. 2018; Richter, & de Lange, 2019; Turk-Browne et. al., 2005) and similar in magnitude to RT benefits elicited by other cognitive factors such as probabilistic attentional cues (Posner, 1980).

We speculate that selective attention may provide a parsimonious explanation for the observed augmented learning in the blocked condition in Experiment 1. Several recent studies show that attentional allocation may proceed in order to maximize learning. For example, observers preferentially attend to stimuli that are not completely predictable or unpredictable (Gottlieb et al., 2013; Kidd et al., 2012; Poli et al., 2020). In other words, their attention is drawn to stimuli that offer maximum information gain. In our experiment, the association between the antedating leading object (A) and the trailing object was learnt during the first training phase. Therefore, participants’ attention may have shifted to the novel blocked (B) leading image during the second training phase, enhancing learning of the association between the blocked leading image and the trailing image. On the other hand, in the control (D) condition, two novel leading objects were presented in the second training phase. In line with overshadowing, these two leading objects may have competed for associative strength with the trailing object and hence their individual predictive power was reduced (Rescorla & Wagner, 1972). In Experiment 2, we aimed to eliminate this attentional effect by applying a backward blocking procedure. Given that the blocked leading object (B) was presented together with a companion leading object (A) in training phase 1, both the companion leading object (A) and blocked leading object (B) were equally familiar and salient in the first phase of the study. As a result, we removed the potentially confounding factors related to novelty and salience, and crucially we provided evidence that backward blocking occurs in statistical learning.

One may wonder whether the present forward and backward blocking experiments provide contradictory results regarding the presence of blocking in statistical learning. Here, it is worth noting that it is more difficult to obtain backward blocking than forward blocking, because more criteria need to be met to observe backward blocking (Van Hamme & Wasserman, 1994). Forward blocking only requires a strong A→X association, which is learned in the elemental training phase, to prevent learning the relationship between cue B and outcome X during the compound training phase. On the other hand, in backward blocking, a strong A→X association learned in the elemental training phase is not enough to observe blocking. In addition to that, the second important condition of backward blocking is that cue A needs to be associated with cue B in order to decrease the associability of cue B in its absence, which is supported by previous studies (Luque et al., 2013; Melchers et al., 2006; Melchers et al., 2004).

By observing backward blocking in Experiment 2, we provide evidence of retrospective revaluation in statistical learning. Such retrospective revaluation cannot be explained by the traditional Rescorla – Wagner model, which assumes that the relevant cue must be present in order to change the associative strength (Kruschke, 2008; Miller & Witnauer, 2016; Rescorla & Wagner, 1972). However, a number of alternative models are able to explain this observation. Van Hamme and Wasserman (1994) proposed a modification of the traditional Rescorla – Wagner model, by allowing an update in the weight of an absent cue if the cue that is associated with the absent cue is present in that trial. Backward blocking can also be explained by a Bayesian generalization of the Rescorla – Wagner model, the Kalman filter (Gershman, 2015; Kruschke, 2008). In sum, our results in Experiment 2 can be explained by both the Van Hamme – Wasserman model and the Kalman filter, both of which claim that learning is based on prediction errors (Kruschke, 2008). At the computational level, this implies that statistical learning may be error-driven. At the implementation level, it supports the view that statistical learning may follow the principles of predictive coding (Hasson, 2017).

Critically, retrospective revaluation may be explained also by the probabilistic contrast model, which does not rely on prediction error (P. Cheng, 1997; P. W. Cheng & Novick, 1992). This model simply calculates how frequently events occur during learning. That is, X appears after either AB or A during training phases, and the probability of X increases after A and in the absence of B. As a result, observers associate A with X. Given that the probabilistic contrast model disregards the order of elemental (i.e., A->X) and compound (i.e., AB->X) training phases (De Houwer & Beckers, 2002), it explains both forward and backward blocking using the same approach. Although our backward blocking results can be explained by the probabilistic contrast model, the model fails to explain our forward blocking results. This supports the importance of the order of training phases in blocking (De Houwer & Beckers, 2002).

Furthermore, it is important to acknowledge that blocking may not arise due to learning deficits, as explained by the models reviewed above, but instead may depend on the failure to express cue – outcome associations at test, as explained by the so-called comparator hypothesis (Miller & Matzel, 1988; Miller & Witnauer, 2016). In other words, retrospective revaluation would not occur because of the increase or decrease in the associative strength between cue and outcome, but rather because of a change in its expression at test. Although we observed backward blocking in statistical learning in Experiment 2, we do not know whether it is because of a learning deficit during training or because of a performance deficit observed at test. Thus, further studies are required to better understand the cause underlying backward blocking in statistical learning.

Crucially, learning regularities is usually incidental rather than intentional in statistical learning paradigms (Batterink et al., 2019; Batterink & Paller, 2017; Henin et al., 2021; Turk-Browne et al., 2005, 2009). However, this can nevertheless result in the development of explicit knowledge of incidentally learned regularities (Batterink & Paller, 2017; Fiser & Aslin, 2002; Turk-Browne et al., 2005). Indeed, testing for explicit knowledge is often used to assess successful incidental learning (Fiser & Aslin, 2002; Turk-Browne et al., 2005). In Experiment 1 and Experiment 2, we observed that people developed some minimal amount of explicit knowledge (on average 56% correct, with chance level of 50%) in the pair recognition test (i.e., how likely the trailing object was given the leading object). The temporal association between leading and trailing object was unknown to participants at the beginning of the experiment and participants were not instructed to learn these associations. Also, participants performed the categorization task at ceiling level (overall above 97%), suggesting that their categorization judgments were not affected by knowledge of the statistical structure between stimuli. Therefore, it appears likely that learning occurred incidentally during the training phases. This is a clear difference between our studies and previous ‘classic’ blocking paradigms where learning occurs intentionally and in the presence of reinforcement (Aggarwal et al., 2020; De Houwer et al., 2002; Kruschke & Blair, 2000; Le Pelley et al., 2005, 2007; Mitchell et al., 2006).

Further, in the context of reinforcement learning, some highlight the key role of inferential reasoning for blocking to occur. Accordingly, learning associations between events does not depend on transitional probabilities but, instead, depends on the observers’ belief about the nature of the relationship between cue and outcome (De Houwer & Beckers, 2002; Waldmann, 2000; Waldmann & Holyoak, 1992). Specifically, the intentional evaluation of causal associations between cues and outcomes (e.g., cue A is the cause of outcome X) appears necessary for forward blocking (De Houwer et al., 2005; De Houwer & Beckers, 2003; Vandorpe et al., 2005) and backward blocking (De Houwer & Beckers, 2002; Waldmann, 2000; Waldmann & Holyoak, 1992). As a result, there is so far evidence that conscious inferential reasoning contributes to backward blocking. To the best of our knowledge, the present study is the first to examine backward blocking in incidental statistical learning. In our experiment, participants were not instructed about any possible relationship between leading and trailing objects, and they learned the associative relationship incidentally. Therefore, our finding supports that conscious inferential reasoning is not required for backward blocking to occur; instead, retrospective revaluation can happen during automatic and incidental statistical learning.

To sum up, we provide evidence for the absence of forward blocking in statistical learning. On the other hand, we provided evidence that backward blocking occurs in statistical learning, where learning developed incidentally and in the absence of rewarding outcomes or feedback. Our results are compatible with the Van Hamme – Wasserman model and Kalman filter, and thus support the idea that statistical learning may be error-driven, similar to reinforcement learning (though see the comparator hypothesis). Most importantly, our results suggest a retrospective revaluation process in statistical learning and thus suggest a link between statistical learning and reinforcement learning.

## Supplementary information 1

### Supplementary text

In addition to the Bayesian analysis reported in the main text, we conducted classic frequentist analyses using R (i.e., ANCOVA of reaction time data in the test phase and one-sample t test and one-way ANOVA of accuracy data in the pair recognition test) for comparability with previous studies and to verify that our conclusions do not depend on the analytical framework employed.

### Analyses of RT data in test phase using ANCOVA

In line with the Bayesian analysis, we first conducted a one-way ANCOVA to determine whether there was a significant difference between reaction times of expected and unexpected trials in the antedating condition, while controlling for the amount of exposure. Secondly, we performed a 2 (Expectation: expected, unexpected) × 2 (Condition: control, blocked) ANCOVA to determine whether there was a significant difference between reaction times of expected and unexpected trials in the control and blocked conditions, while controlling for the amount of exposure. In line with the primary analysis, the contrasts of these factors were coded as successive difference contrasts.

In Experiment 1, the main effect of Expectation was significant in the antedating condition after controlling for Exposure (F(1, 792) = 106e+4, p < 0.001, partial η^2^ = 0.93). Across control and blocked trials, the main effect of Expectation (F(1, 1577) = 4.33, p = 0.04, partial η^2^ = 2.74e-3) was significant, but the main effect of Condition (F(1, 1577) = 0. 62, p = 0.43, partial η^2^ = 3.93e-4) was not significant. The Expectation × Condition interaction was not significant (F(1, 1577) = 0.20, p = 0.65, partial η^2^ = 1.28e-4). Overall, in Experiment 1, the ANCOVA analysis confirmed the results of the Bayesian mixed effect model analysis: in the antedating condition, we found successful learning of repeated stimulus pairs; crucially, we found no blocking effect for incidentally learned stimulus pairs.

In Experiment 2, the main effect of Expectation was significant in the companion condition after controlling for Exposure (F(1, 795) = 12143, p < 0.001, partial η^2^ = 0.94). Across control and blocked trials, the main effect of Expectation (F(1, 1582) = 0.01, p = 0.93, partial η^2^ = 5.57e-6) and the main effect of Condition (F(1, 1582) = 0. 02, p = 0.89, partial η^2^ = 1.14e-5) were not significant. The Expectation × Condition interaction was not significant (F(1, 1582) = 1.98, p = 0.16, partial η^2^

= 1.25e-3). Overall, in Experiment 2, the ANCOVA analysis did not confirm the results of the Bayesian mixed effect model analysis: in the companion condition, we found successful learning of repeated stimulus pairs.

### Analyses of accuracy data in pair recognition test using ANOVA and t-test

In line with the Bayesian analysis, we first conducted a one-sample t-test to determine whether the level of accuracy was above chance level in each condition. Secondly, we performed a one-way (Condition: control – blocked) ANOVA to test for the blocking effect.

In Experiment 1, the level of accuracy was above chance level in the antedating (t(49) = 6.37, p < 0.001, Cohen’s d = 0.90), blocked (t(49) = 4.60, p < 0.001, Cohen’s d = 0.65) and control (t(49) = 5.48, p < 0.001, Cohen’s d = 0.78) conditions. Furthermore, the main effect of Condition (F(1, 98) = 1.01, p = 0.31, partial η^2^ = 0.01) was not significant, indicating the absence of forward blocking effect for the explicit knowledge of incidentally learned associations. Overall, in Experiment 1, the t test and ANOVA analysis confirmed the results of the Bayesian mixed effect model analysis reported in the main text: we found clear explicit knowledge of incidentally learned associations in each condition and no blocking effect for such explicit knowledge.

In Experiment 2, the level of accuracy was above chance level in the companion (t(49) = 3.64, p = 0.00, Cohen’s d = 0.51) condition. Overall, in Experiment 2, the t test analysis confirmed the results of the Bayesian mixed effect model analysis reported in the main text: we found clear explicit knowledge of incidentally learned associations in companion condition but not in blocked and control conditions.

### Supplementary tables

**Table S1.** Posterior fixed effects of the post-hoc model of control condition on reaction times in Experiment 1. Estimate, estimation error, lower/upper limit of 95% profile credible intervals.

| Predictors | *Estimate* | *Est. Error* | *CI (95%)* |
| --- | --- | --- | --- |
| Intercept | 491.09 | 9.73 | 472.16 – 510.23 |
| Expectation | 4.36 | 2.59 | -0.73 – 9.51 |
| Exposure | -30.03 | 4.33 | -38.34 – -21.45 |
| Expectation ×<br>Exposure | -2.02 | 3.71 | -9.26 – 5.21 |

**Table S2.**
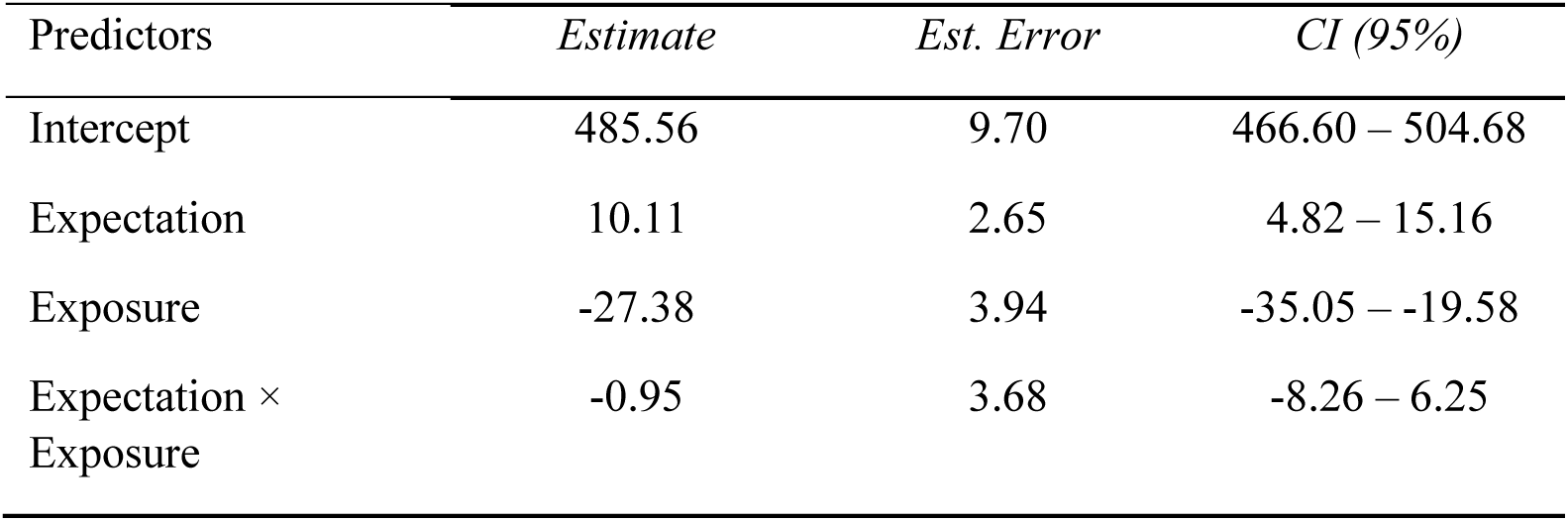
Posterior fixed effects of the post-hoc model of blocked condition on reaction times in Experiment 1. Estimate, estimation error, lower/upper limit of 95% profile credible intervals.

**Table S3.** Posterior fixed effects of the post-hoc model of control condition on reaction times in Experiment 2. Estimate, estimation error, lower/upper limit of 95% profile credible intervals.

| Predictors | *Estimate* | *Est. Error* | *CI (95%)* |
| --- | --- | --- | --- |
| Intercept | 500.48 | 9.47 | 481.97 – 519.28 |
| Expectation | 8.44 | 2.46 | 3.60 – 13.29 |
| Exposure | -22.24 | 4.70 | -31.37 – -12.93 |
| Expectation ×<br>Exposure | 0.38 | 3.57 | -6.43 – 7.36 |

**Table S4.**
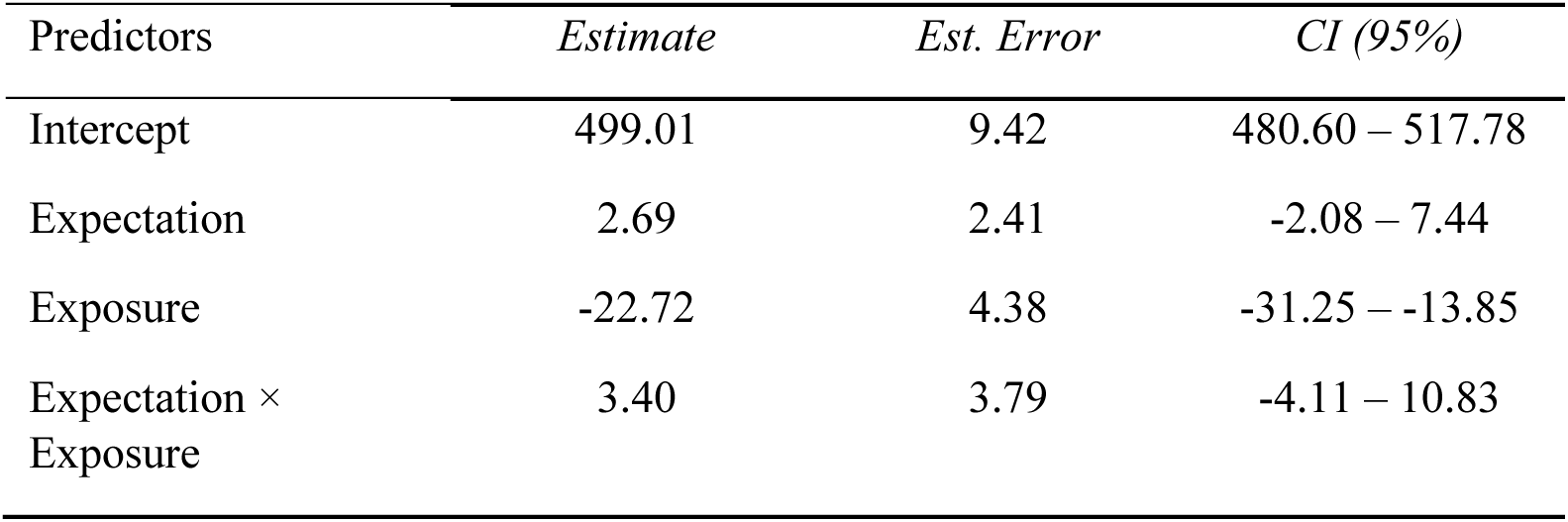
Posterior fixed effects of the post-hoc model of blocked condition on reaction times in Experiment 2. Estimate, estimation error, lower/upper limit of 95% profile credible interval.

## Supplementary information 2

## Supplementary Experiment 1

## Method

### Participants

The experiment was performed online using the Gorilla platform (Anwyl-Irvine et al., 2020), and participants were recruited through the Prolific platform (https://www.prolific.co/). 148 participants performed the experiment. 47 of them were excluded before they finished the experiment based on a priori exclusion criteria (see section ‘Exclusion and inclusion criteria’), and one participant was excluded from the final data analysis due to excessively slow responses (RTs above 3 times the mean absolute deviation [MAD] from the group mean). As a result, one hundred participants (37 females; mean age 24.49, range 18-40 years) were included in the data analysis. This final number of included participants was preregistered based on previous research (Richter & de Lange, 2019; Schmidt & De Houwer, 2019) considering that online data would be noisier and, therefore, a larger number of participants would be required to maintain the same statistical power. The pre-selected sample size yielded 84% power to detect a small sized (Cohen’s *d*_z_ = 0.3) effect (α = 0.05).

### Experimental design

The design and procedure of Supplementary Experiment 1 was identical in all respects to Experiment 1 (see Figure S1a). In line with the traditional blocking paradigm Supplementary Experiment 1 comprised two training phases (training phase 1 and training phase 2) and a test phase. During the two training phases, leading object were perfectly predictive of their respective trailing object (i.e. P(trailing | leading = 1) ; see Figure S1b). Yet during the final test stage Expected and unexpected object pairs were presented equally often to prevent any learning (see Figure S1c). The main difference between Supplementary Experiment 1 and Experiment 1 is related to the type of leading stimuli and stimulus location. Leading stimulus was either a geometric shape or an everyday object. If the antedating leading stimulus was an object, then the blocked leading stimulus was a shape or vice versa. Both leading and trailing stimuli were presented at the center of the screen.

### Data analysis

The data analysis of Supplementary Experiment 1 was identical in all respects to Experiment 1, except for the priors and the additional analysis of RT data *split by stimulus type in test phase*.

The models were constructed using weakly informative priors. The prior distributions for the effects of interest were Gaussian distributions with zero mean and standard deviation adjusted to expected effect sizes: 50 for Expectation, 70 for Condition and 30 for Exposure, 70 for the interaction between Expectation and Condition. Further details and the complete model parametrization can be found in the R codes provided on the Donders Repository.

#### Analysis of RT data split by stimulus type in test phase

We conducted a follow-up analysis that tested for the effect of the type of leading stimulus (shape / object). We reasoned that leading object stimuli may have attracted more attention than leading shape stimuli, given that they were visually more salient than the surrounding grey shapes, and their category was task-relevant, as the task required object categorization on the trailing image. Given that associative learning depends on attention (Kruschke, 2001; Pacton & Perruchet, 2008), it was therefore conceivable that leading objects, rather than shapes, developed a stronger temporal association with trailing objects. We fit the model of antedating condition and the model of blocked and control conditions as described above, but with the inclusion of leading Stimulus Type (shape / object) as additional fixed factor. The model included a full random effect structure (i.e., a random intercept and slopes for all within-participant effects). If the posterior credible intervals of the interaction effects between Expectation and leading Stimulus Type did not overlap with zero, we run separate models for shapes and objects respectively, in order to test for the blocking effect for each stimulus type. The models were constructed using weakly informative priors centered at zero. All other analysis settings were as specified above.

### Results

#### Analysis of RT data in test phase

First, we compared the reaction times of expected and unexpected trials in the antedating condition to test whether repeated exposure to leading-trailing pairs led to learning their temporal association (see Table S1). We observed faster reaction times in expected (493 ms) than unexpected (508 ms) trials (b = 11.23, CI = [6.80, 15.59], Cohen’s *d*_z_ = 0.54, see Figure 1d), indicating successful learning of stimulus transition probabilities and the consequent behavioral benefit of expectation in terms of response speed. In addition, we tested whether this behavioral benefit remained stable during the test phase or dwindled, as would be expected by extinction. In line with the latter, we observed an interaction effect between Expectation and Exposure (b = -9.28, CI = [-15.26, -3.38]), indicating that learning showed rapid extinction (expectation effect for run 1: 22 ms, run 2: 9 ms, run 3: 6 ms; see Figure S1e).

Next, we moved to our main question and tested for the presence of blocking (see Table S2 and Figure 1f). The reaction time difference between unexpected and expected trials was not different between control (11 ms) and blocked (12 ms) conditions (b = 1.85, CI = [-3.95, 7.51], Cohen’s *d*_z_ = -0.04, see Figure S1d). This pattern of results presents suggest for the absence of blocking. There was also no difference in how the reaction time benefit for expected items behaved over time (b = -2.29, CI = [-11.17, 6.13]; expectation effect in blocked condition for run 1: 13 ms, run 2: 4 ms, run 3: 12 ms; expectation effect in control condition for run 1: 18 ms, run 2: 10 ms, run 3: 7 ms; see Figure S1e).

#### Analyses of RT data split by stimulus type in test phase

In a follow-up analysis, we tested whether the type of leading stimulus (shape / object) affected statistical learning. In the antedating condition (see Table S3), the reaction time difference between unexpected and expected trials was larger for leading object (20 ms) compared to leading shape (9 ms) trials according to the posterior CI (b = -10.00, CI = [-18.57, -1.48]), which indicated that object-object associations were somewhat stronger than shape-object associations. While the difference in RT was larger for object-object associations than shape-object associations, separate follow-up models showed that the reaction time difference was significant when the leading stimulus was an object (b = 15.19, CI = [7.98, 22.46], see Table S1 and Figure S2a-e), and it was still significant (b = 5.44, CI = [0.83, 10.05], Table S2 and see Figure S2b-f).

Across blocked and control conditions (see Table S4), the reaction time difference between unexpected and expected trials was also larger when the leading stimulus was an object (18 ms for B, 27 ms for D) compared to a shape (0 ms for B, 1 ms for D) (b = 18.40, CI = [11.52, 25.41]). Separate follow-up models showed that reaction times were faster in expected trials than in unexpected trials when the leading stimulus was an object (RT difference = 18 ms in blocked condition and 27 ms in control condition; b = 18.73, CI = [12.83, 24.5], see Table S3 and Figure S2a-e). This was not the case when the leading stimulus was a shape (RT difference = 0 ms in blocked condition and 1 ms in control condition; b = 0.11, CI = [-3.27, 3.44], Table S4 and see Figure S2b-f). Overall, the data suggest that shape – object associations could be learnt, but to a lesser extent than object – object associations. In particular, shape – object associations could be learnt only if a leading shape in isolation was followed by a trailing object (i.e., in the antedating condition), but not when the leading shape was concurrently paired with a leading object (in a compound stimulus) and then followed by the trailing object (i.e., in the blocked and control conditions). This pattern of results fits our prediction that leading objects attract more attention than shapes, given that they were visually more salient, and their category was task-relevant. As associative learning depends on attention (Kruschke, 2001; Pacton & Perruchet, 2008), this may have hampered associative learning between leading shapes and trailing objects. In other words, we found cue competition among the leading shape and object in the forms of overshadowing (Boddez et al., 2014; Pavlov, 1927; Schmidt & De Houwer, 2019), with the leading shape being overshadowed by the leading object. Finally, there was evidence for the absence of an interaction between Expectation, Condition and leading Stimulus Type (b = 4.09, CI = [-6.18, 15.80]), indicating that the absence of blocking did not depend on leading Stimulus Type.

#### Analyses of accuracy data in pair recognition test

Participants showed slightly above chance-level performance in indicating whether the trailing object was likely or unlikely given the leading stimulus in the antedating (proportion correct = 58%; b = 0.32, CI = [0.23, 0.42]), blocked (proportion correct = 54%; b = 0.16, CI = [0.09, 0.24]) and control (proportion correct = 53%; b = 0.12, CI = [0.04, 0.19]) conditions (see Figure S1g). Response errors did not differ between the blocked and control conditions (b = -0.05, CI = [-0.15, 0.05]), indicating no blocking for the explicit knowledge of incidentally learned associations.

### Supplementary Experiment 2

Supplementary Experiment 1 showed that the type of leading stimulus critically influenced statistical learning. Antedating and control leading shapes got less strongly associated with the trailing object than antedating and control leading objects. Moreover, blocked and control leading shapes could not compete with the concurrent leading objects for associative strength because they attracted less attention. This imbalance between shapes and objects may provide an alternative explanation for the lack of blocking that we observed. Therefore, in Supplementary Experiment 2

we made one modification to our paradigm and only presented objects as leading and trailing stimuli to remove any potential difference in attention between different leading stimuli, which might finally result in a blocking effect.

## Method

### Participants

The experiment was performed online by using the Gorilla platform (Anwyl-Irvine et al., 2020), and participants were recruited through the Prolific platform (https://www.prolific.co/). 81 participants performed the experiment. 27 of them were excluded before they finished the experiment based on a priori exclusion criteria (see section ‘Exclusion and inclusion criteria’ above). Four extra participants were excluded from the final data analysis: two showed accuracy below 50% chance level in test phase; two showed overall excessively slow responses (RTs above 3 MAD from the group mean). As a result, fifty participants (16 females; mean age 23.90, range 18-34 years) were included in the data analysis, as preregistered. This final number of included participants was derived from the following a priori power calculation: we aimed for 90% power to detect the effect size of Cohen’s *d*_z_ = 0.5 derived in the antedating leading object condition of Experiment 1 (α = 0.05).

All participants had normal or corrected to normal vision, normal hearing and no history of neurological or psychiatric conditions. They provided written informed consent and received financial reimbursement (8 euros per hour) for their participation in the experiment. The study followed the guidelines for ethical treatment of research participants by CMO 2014/288 region Arnhem-Nijmegen, The Netherlands.

### Experimental design

The design and procedure of Supplementary Experiment 2 was identical in all respects to Supplementary Experiment 1, apart from the type of leading stimuli and their location. Both leading and trailing stimuli were everyday objects. Leading and trailing objects were randomly presented on the left or right side of the central fixation point. Stimuli position (left / right) was counterbalanced with respect to Expectation (expected / unexpected) and Condition (antedating / blocked / control). In other words, leading and trailing objects appeared equally often on the left or right side of the central fixation point across trials. As a result, the expectation manipulation did not depend on spatial position. In addition, both hemi-fields were equally task-relevant, which fostered participants’ attention to both sides.

### Data analysis

The data analysis of Supplementary Experiment 2 was identical in all respects to Supplementary Experiment 1, except for omitting the factor Stimulus Type because this experiment featured only object stimuli (see Figure S3a).

### Results

#### Analyses of RT data in test phase

First, we compared the reaction times of expected and unexpected trials in the antedating condition (see Table S5). We observed that reaction times for expected (503 ms) and unexpected (510 ms) trials, although showing a qualitative pattern similar to Experiment 1, were not significantly different from each other (b = 4.95, CI = [-0.07, 9.96], Cohen’s *d*_z_ = 0.31, see Figure S3b). Therefore, unlike Experiment 1, our data do not provide robust support for learning of the conditional probabilities in condition A. There was however some statistical support for extinction, as the reaction time difference between expected and unexpected trials tended to decrease as the exposure increased (b = -8.17, CI = [-15.39, -0.91]) (expectation effect for run 1: 17 ms, run 2: 6 ms, run 3: 0 ms; see Figure S3c).

Next, we moved to our main question and compared reaction time differences between expected and unexpected stimulus pairs between B and C (see Table S6 and Figure S3d). The reaction time difference between unexpected and expected trials was not statistically different between control (8 ms) and blocked (1 ms) conditions (b = 3.34, CI = [-3.11, 9.85], Cohen’s *d*_z_ = 0.24, see Figure S3b). Moreover, extinction was not different between B and C (b = 0.37, CI = [-9.60, 10.22]; expectation effect in blocked condition for run 1: 6 ms, run 2: -2, run 3: 0 ms; expectation effect in control condition for run 1: 11 ms, run 2: 4 ms, run 3: 5 ms; see Figure S3c).

#### Analysis of accuracy data in pair recognition test

Participants were not able to indicate above chance level whether the trailing object was likely or unlikely given the leading object in the antedating (proportion correct = 50%; b = 0, CI = [-0.15, 0.14]), blocked (proportion correct = 49%; b = -0.05, CI = [-0.17, 0.07]) or control (proportion correct = 50%; b = 0, CI = [-0.13, 0.14]) conditions (see Figure S3e).

## Supplementary figures

**Figure S1.**
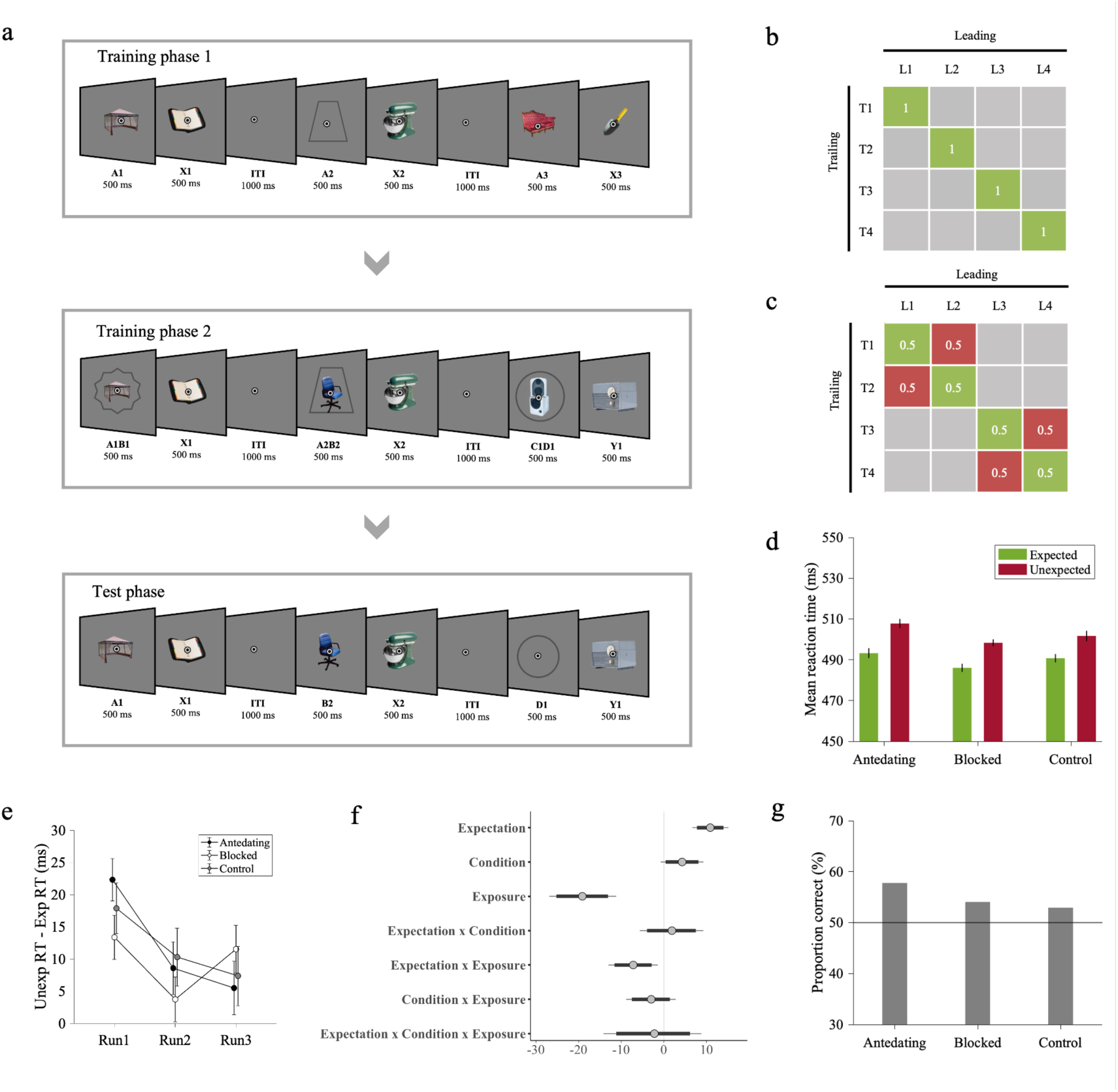
*Experimental procedure and results of* Supplementary Experiment 1. *Note.* (a) Experiment 1 comprised two training phases (training phase 1 and training phase 2) and a test phase. On every trial throughout the experiment, participants saw a pair of consecutively presented stimuli, i.e., a leading image followed by a trailing image. In training phase 1, the antedating leading stimulus (i.e., A), which could be either a shape or object, was followed by a specific trailing object. In training phase 2, a novel blocked leading stimulus (i.e., B) was presented in compound, along with the antedating (A) leading stimulus (i.e., AB), and followed by the same trailing object from the antedating stimulus in training phase 1. In addition, we introduced novel control compound leading (i.e., CD) and trailing (i.e., Y) stimuli. In the test phase, antedating, blocked or control leading stimuli were followed by the associated (expected) or not associated (unexpected) trailing object. There were four different object pairs for AB→X and CD→Y. Throughout the experiment, participants performed a categorization task on the trailing object. They reported, as fast as possible, whether the trailing object was electronic or non-electronic. (b) Statistical regularities depicted as image transition matrix with stimuli pairs in training phase 1 and training phase 2. Ls represent leading stimuli, and Ts represent trailing stimuli. There were 16 different leading objects and 8 different trailing objects coming from four different AB→X and CD→Y pairs. (c) Statistical regularities depicted as image transition matrix with stimuli pairs in test phase. Green cells represent expected pairs, and red cells represent unexpected pairs. (d) Across participants’ mean reaction times as a function of Expectation (expected / unexpected) and Condition (antedating / blocked / control). Participants responded faster to expected than unexpected trailing objects in each condition. There was no difference between blocked and control conditions. (e) Across participants’ mean reaction time difference between expected and unexpected trials as a function of time. Please note that we split data into successive runs for visualization purposes only; data analysis was performed with number of trials as a continuous fixed factor (Exposure). Associations were rapidly extinguished during the test phase. Extinction was not different between conditions. (f) Posterior coefficient estimates of effects of the model jointly analyzing blocked and control conditions with error bars representing 95% confidence intervals. Estimates indicate significant results when they do not overlap with zero. (g) Across participants’ proportion correct responses in pair recognition test. Participants showed slightly above chance-level performance in indicating whether the trailing object was likely or unlikely given the leading stimulus in all conditions.

**Figure S2.**
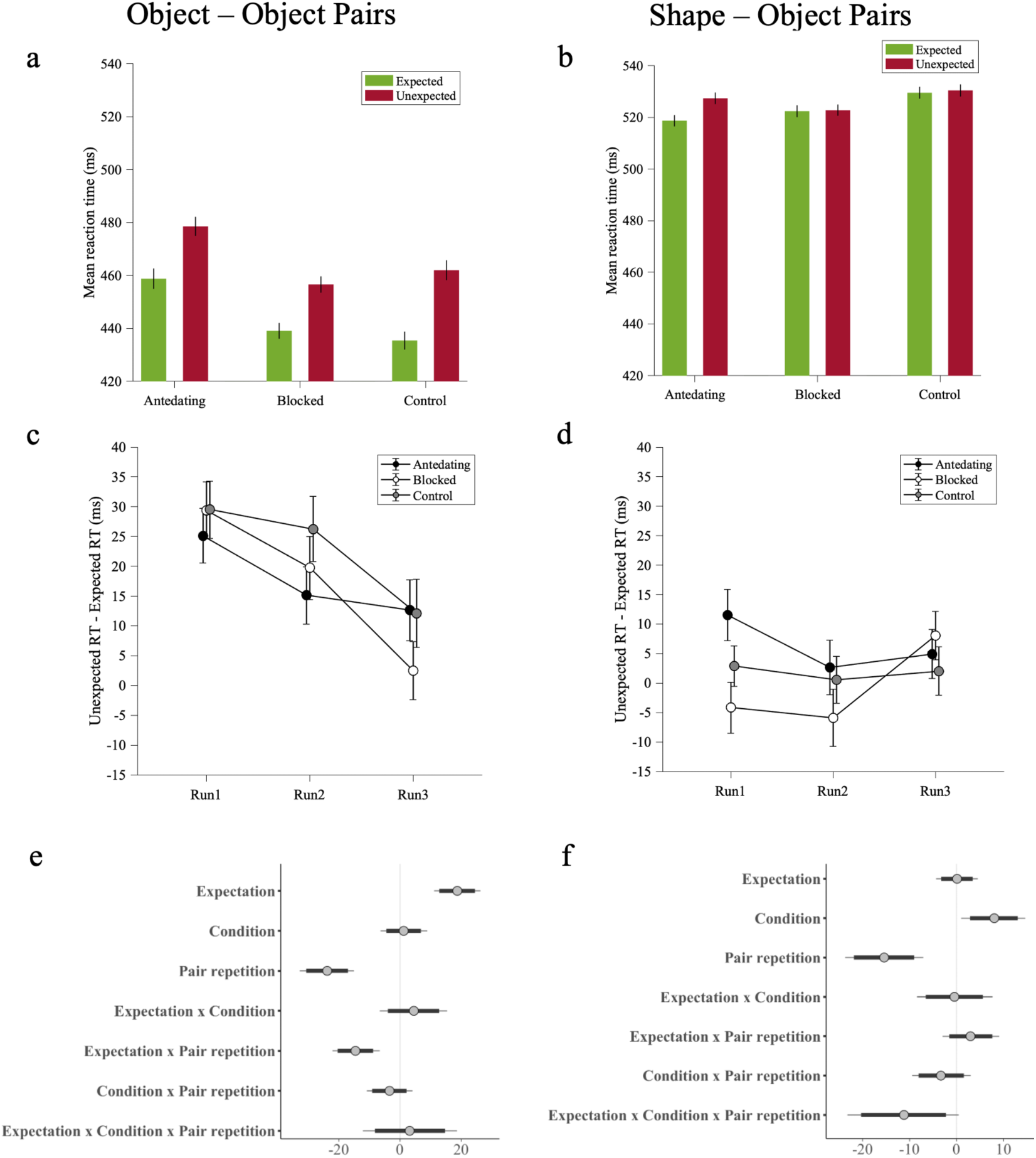
*Results of* Supplementary Experiment 1 *as a function of Stimulus Type*. *Note.* (a-b) Across participants’ mean reaction times as a function of Expectation (expected / unexpected) and Condition (antedating / blocked / control) in leading objects (a) and leading shapes (b). The difference between expected and unexpected reaction times was larger for stimulus pairs with leading objects, compared to leading shapes. (c-d) Across participants’ mean reaction time difference between expected and unexpected trials as a function of time in leading objects (c) and leading shapes (d). The decrease in reaction time difference between expected and unexpected trials over exposure showed rapid extinction in learning only in leading objects. (e-f) Posterior coefficient estimates of effects of the model jointly analyzing blocked and control conditions with error bars representing 95% confidence intervals in leading objects (e) and leading shapes (f). Estimates indicate significant results when they do not overlap with zero.

**Figure S3.**
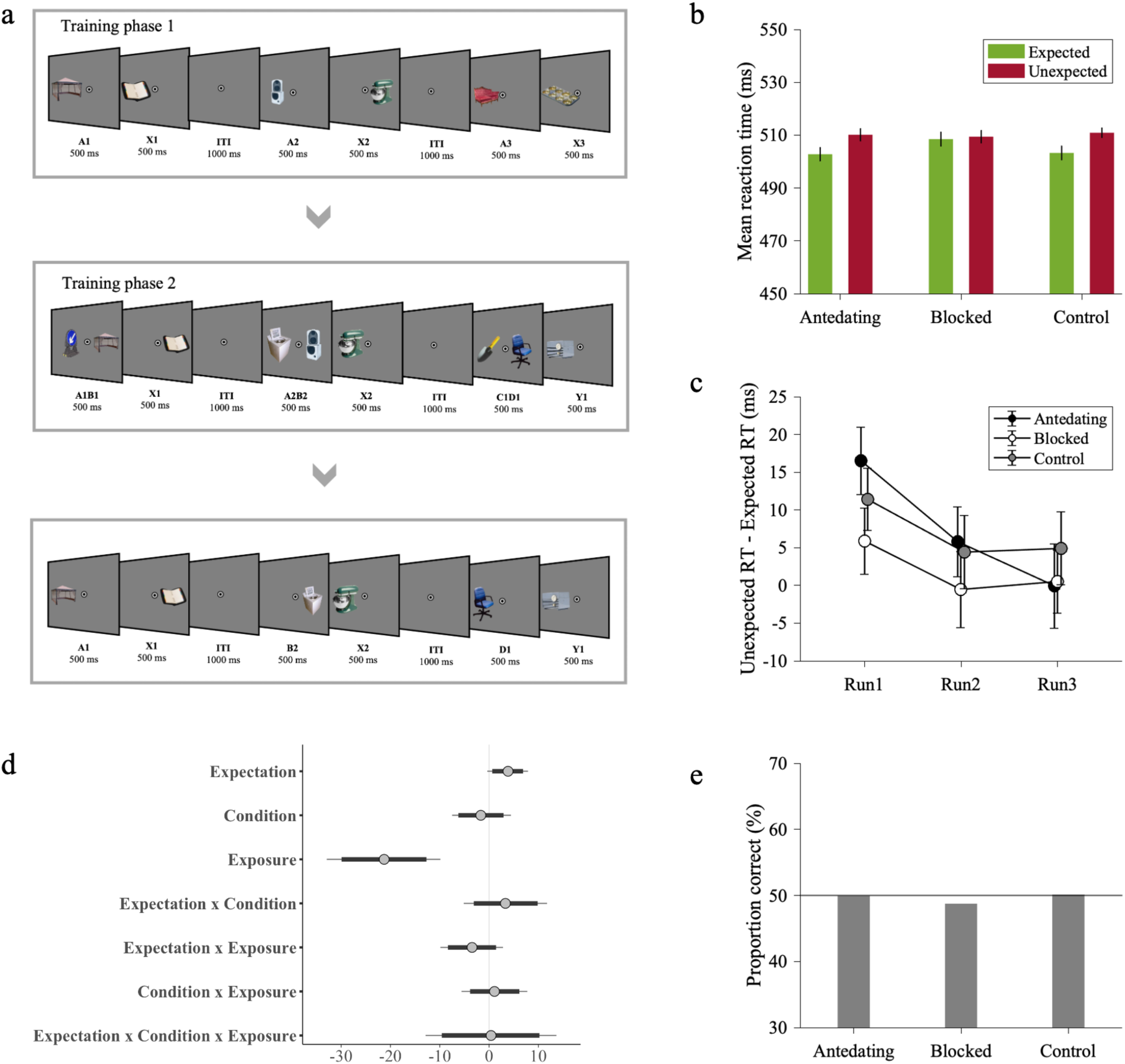
*Experimental procedure and results of* Supplementary Experiment 2. *Note.* (a) The design and procedure of experiment 2 was identical in all respects to experiment 1, apart from the fact that the leading stimulus was an object presented in the left or right side of the fixation point, and it was followed by the trailing object presented in the left or right side of the fixation point. (b) Across participants’ mean reaction times as a function of Expectation (expected / unexpected) and Condition (antedating / blocked / control). Reaction times were faster to expected than unexpected trailing objects in blocked and control conditions. There was no difference between blocked and control condition in terms of reaction time difference between expected and unexpected trials, providing evidence for the absence of blocking effect. (c) Across participants’ mean reaction time difference between expected and unexpected trials as a function of time. The decrease in reaction time difference between expected and unexpected trials over exposure showed rapid extinction in learning antedating condition. (d) Posterior coefficient estimates of effects of the model jointly analyzing blocked and control conditions with error bars representing 95% confidence intervals. Estimates indicate significant results when they do not overlap with zero. (e) Across participants’ proportion correct responses in pair recognition test. Participants were not able to indicate above chance level whether the trailing object was likely or unlikely given the leading object in all conditions.

## Supplementary tables

**Table S1.** Posterior fixed effects of the model of antedating condition on reaction times in Experiment 1. Estimate, estimation error, lower/upper limit of 95% profile credible intervals.

| Predictors | *Estimate* | *Est. Error* | *CI (95%)* |
| --- | --- | --- | --- |
| Intercept | 502.44 | 8.42 | 485.21 – 518.66 |
| Expectation | 11.23 | 2.25 | 6.80 – 15.59 |
| Exposure | -15.14 | 3.51 | -22.08 – -8.19 |
| Expectation ×<br>Exposure | -9.28 | 3.01 | -15.26 – -3.38 |

**Table S2.** Posterior fixed effects of the model of blocked and control conditions on reaction times in Experiment 1. Estimate, estimation error, lower/upper limit of 95% profile credible intervals.

| Predictors | *Estimate* | *Est. Error* | *CI (95%)* |
| --- | --- | --- | --- |
| Intercept | 494.45 | 8.34 | 478.02 – 510.93 |
| Expectation | 10.88 | 1.6 | 7.76 – 13.98 |
| Condition | 4.30 | 1.95 | 0.38 – 8.10 |
| Exposure | -19.10 | 3.08 | -25.19 – -13.08 |
| Expectation ×<br>Condition | 1.85 | 2.91 | -3.95 – 7.51 |
| Expectation ×<br>Exposure | -7.19 | 2.24 | -11.61 – -2.87 |
| Condition ×<br>Exposure | -3.00 | 2.26 | -7.49 – 1.40 |
| Expectation ×<br>Condition ×<br>Exposure | -2.29 | 4.48 | -11.17 – 6.13 |

**Table S3.** Posterior fixed effects of the model of antedating condition on reaction times split by stimulus type in Experiment 1. Estimate, estimation error, lower/upper limit of 95% profile credible intervals.

| Predictors | *Estimate* | *Est. Error* | *CI (95%)* |
| --- | --- | --- | --- |
| Intercept | 502.27 | 8.42 | 485.68 – 518.72 |
| Expectation | 10.37 | 2.18 | 6.15 – 14.62 |
| Leading stimulus type | 56.95 | 5.6 | 46.13 – 67.99 |
| Exposure | -15.35 | 3.52 | -22.36 – -8.31 |
| Expectation × Leading stimulus type | -10.00 | 4.37 | -18.57 – -1.48 |
| Expectation × Exposure | -7.26 | 2.61 | -12.36 – -2.18 |
| Leading stimulus type × Exposure | 10.55 | 3.81 | 3.02 – 18.15 |
| Expectation × Leading stimulus type × Exposure | 1.12 | 5.38 | -9.32 – 11.81 |

**Table S4.**
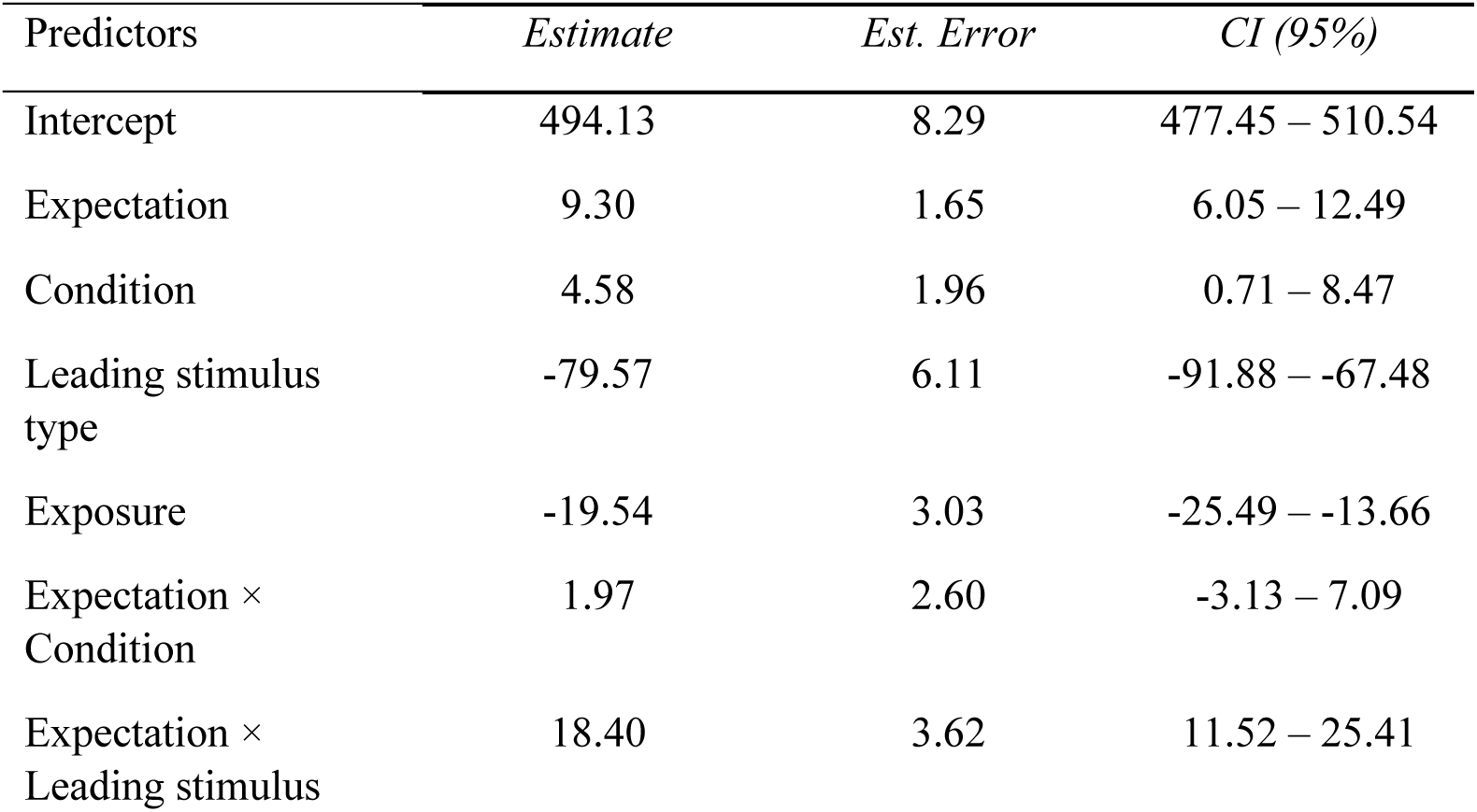
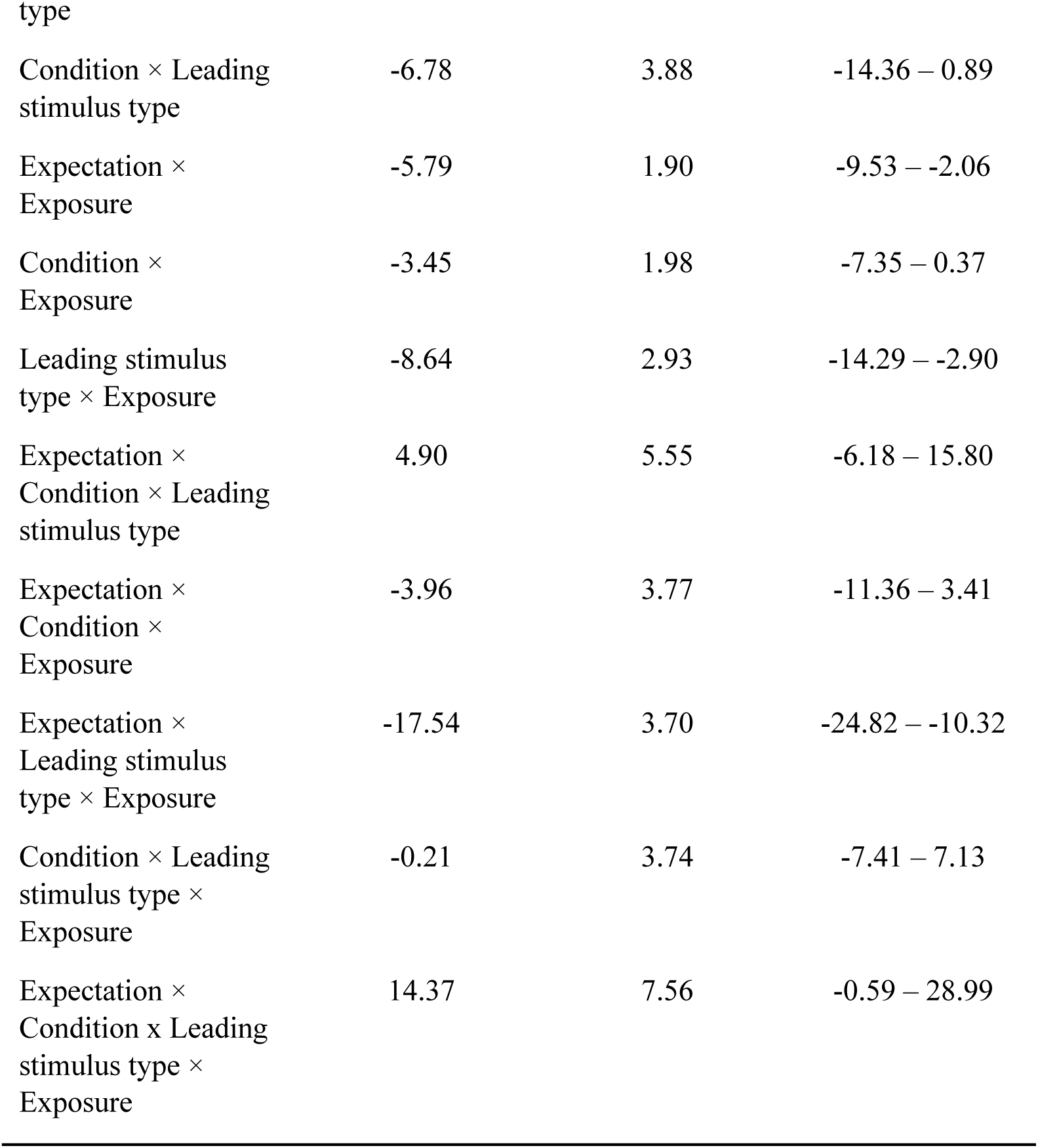
Posterior fixed effects of the model of blocked and control conditions on reaction times split by stimulus type in Experiment 1. Estimate, estimation error, lower/upper limit of 95% profile credible intervals.

| Predictors | *Estimate* | *Est. Error* | *CI (95%)* |
| --- | --- | --- | --- |
| Intercept | 494.13 | 8.29 | 477.45 – 510.54 |
| Expectation | 9.30 | 1.65 | 6.05 – 12.49 |
| Condition | 4.58 | 1.96 | 0.71 – 8.47 |
| Leading stimulus type | -79.57 | 6.11 | -91.88 – -67.48 |
| Exposure | -19.54 | 3.03 | -25.49 – -13.66 |
| Expectation × Condition | 1.97 | 2.60 | -3.13 – 7.09 |
| Expectation × Leading stimulus type | 18.40 | 3.62 | 11.52 – 25.41 |
| Condition × Leading stimulus type | -6.78 | 3.88 | -14.36 – 0.89 |
| Expectation × Exposure | -5.79 | 1.90 | -9.53 – -2.06 |
| Condition × Exposure | -3.45 | 1.98 | -7.35 – 0.37 |
| Leading stimulus type × Exposure | -8.64 | 2.93 | -14.29 – -2.90 |
| Expectation × Condition × Leading stimulus type | 4.90 | 5.55 | -6.18 – 15.80 |
| Expectation × Condition × Exposure | -3.96 | 3.77 | -11.36 – 3.41 |
| Expectation × Leading stimulus type × Exposure | -17.54 | 3.70 | -24.82 – -10.32 |
| Condition × Leading stimulus type × Exposure | -0.21 | 3.74 | -7.41 – 7.13 |
| Expectation × Condition × Leading stimulus type × Exposure | 14.37 | 7.56 | -0.59 – 28.99 |

**Table S5.** Posterior fixed effects of the model of antedating condition on reaction times in Experiment 2. Estimate, estimation error, lower/upper limit of 95% profile credible intervals.

| Predictors | *Estimate* | *Est. Error* | *CI (95%)* |
| --- | --- | --- | --- |
| Intercept | 512.83 | 18.97 | 475.52 – 549.47 |
| Expectation | 4.95 | 2.51 | -0.07 – 9.96 |
| Exposure | -18.40 | 4.29 | -26.74 – -10.02 |
| Expectation × Exposure | -8.17 | 3.70 | -15.39 – -0.91 |

**Table S6.** Posterior fixed effects of the model of blocked and control conditions on reaction times in Experiment 2. Estimate, estimation error, lower/upper limit of 95% profile credible intervals.

| Predictors | *Estimate* | *Est. Error* | *CI (95%)* |
| --- | --- | --- | --- |
| Intercept | 515.30 | 19.42 | 476.89 – 533.55 |
| Expectation | 3.82 | 1.61 | 0.64 – 6.90 |
| Condition | -1.68 | 2.33 | -6.23 – 2.94 |
| Exposure | -21.29 | 4.38 | -29.92 – 12.70 |
| Expectation ×<br>Condition | 3.34 | 3.28 | -3.11 – 9.85 |
| Expectation ×<br>Exposure | -3.47 | 2.51 | -8.35 – 1.42 |
| Condition ×<br>Exposure | 1.12 | 2.58 | -3.86 – 6.15 |
| Expectation ×<br>Condition ×<br>Exposure | 0.37 | 5.08 | -9.60 – 10.22 |

